# Abundance and Microbial Diversity from Surface to Deep Water Layers Over the Rio Grande Rise, South Atlantic

**DOI:** 10.1101/2021.04.22.441028

**Authors:** Juliana Correa Neiva Ferreira, Natascha M. Bergo, Pedro M. Tura, Mateus Gustavo Chuqui, Frederico P. Brandini, Luigi Jovane, Vivian H. Pellizari

## Abstract

Marine microbes control the flux of matter and energy essential for life in the oceans. Until now, the distribution and diversity of planktonic microorganisms above Fe-Mn crusts has received relatively little attention. Future mining\dredging of these minerals is predicted to affect microbial diversity and functioning in the deep sea. Here, we studied the ecology of planktonic microbes among pelagic environments of an Fe-Mn deposit region, at Rio Grande Rise, Southwestern Atlantic Ocean. We investigated microbial community composition using high-throughput sequencing of 16S rRNA genes and their abundance estimated by flow cytometry. Our results showed that the majority of picoplanktonic was found in epi- and mesopelagic waters, corresponding to the Tropical Water and South Atlantic Central Water. Bacterial and archaeal groups related to phototrophy, heterotrophy and chemosynthesis, such as *Synechococcales*, Sar11 (Proteobacteria) and Nitrosopumilales (Thaumarchaeota) were the main representatives of the pelagic microbial community. Additionally, we detected abundant assemblages involved in biodegradation of marine organic matter and iron oxidation at deep waters, i.e., *Pseudoalteromonas* and *Alteromonas*. No differences were observed in microbial community alpha diversity. However, we detected differences in community structure between water masses, suggesting that changes in an environmental setting (i.e. nutrient availability or circulation) play a significant role in structuring the pelagic zones, also affecting the meso- and bathypelagic microbiome.

**Highlights:**

- Rio Grande Rise pelagic microbiome
- Picoplankton carbon biomass partitioning through pelagic zones
- Unique SAR11 Clade I oligotype in the shallowest Tropical Water
- Higher number of shared oligotypes between deepest water masses
- Nitrogen, carbon and sulfur may be important contributors for the pelagic microbiome

## INTRODUCTION

The world’s oceans are an enormous pool of diverse microscopic life forms, that play a vital role in food chains and nutrient cycling (Teeling *et al*., 2012; Hogle *et al*., 2016). Marine picophytoplankton are responsible for almost half of the primary production on the planet, in particular, the genera *Synechococcus* and *Prochlorococcus* (Azam & Malfatti, 2007), capable of adsorbing trace metals (e.g., Mn, Fe, Ni, Cu, Zn, and Cd) from seawater for photosynthetic carbon fixation (Morel *et al*., 2020). Most of the particulate organic matter produced by the phytoplankton in the euphotic zone is excreted as dissolved organic matter which is remineralized by heterotrophic Bacteria and Archaea (Azam & Malfatti, 2007). A small fraction of the particulate organic carbon from primary producers is exported to deeper waters (Briggs *et al*., 2020), as the primary source of energy in and below the meso- and bathypelagic zones. This is particularly relevant to seamount and rise habitats and possibly those associated with ferromanganese (Fe-Mn) deposits due to their proximity to the productive zone above and the hydrodynamic of these regions that may increase the water residence (Genin *et al*., 2007) and microbial processes above seamounts (Mendonça *et al*., 2012; Lemos *et al*., 2018; Giljan *et al*., 2020).

The transport of organic matter produced in the euphotic zone to the benthic ecosystems adjacent to Fe-Mn deposits is relatively low (Smith *et al*., 2008; Wedding *et al*., 2013). Yet, it is an important source of bioactive metals (Smith *et al*., 2008; Wigham *et al*., 2003) and has a strong influence on benthic organisms and community diversity. Besides trace elements for enzyme cofactors, which catalyze key steps in respiration, nitrogen fixation, and photosynthesis, marine phytoplankton and heterotrophic bacterioplankton also require macronutrients for structural components such as proteins and membranes (Butler, 1998; Morel *et al*., 2003).

Equally important, future Fe-Mn crust mining from seamounts is expected to physically alter the seafloor (Rolinski *et al*., 2001), producing a plume of waste material and impacting circulation, chemical characteristics and redox potential of the adjacent water column (Orcutt *et al*., 2020). These changes may potentially alter the vertical transport of organic matter (Langenheder *et al*., 2010) and, consequently, the microbial contribution for the formation of Fe-Mn crusts (Orcutt *et al*., 2020). Although planktonic Bacteria and Archaea communities can rapidly respond to these environmental disturbances (Allison & Martiny, 2008), Orcutt *et al*. (2020) recommend that microbial diversity, biomass, and biogeochemical contributions need to be considered in environmental impact assessments of deep-sea mining.

The Rio Grande Rise (RGR) is a wide elevation (~150,000 km^2^) located in the southwestern portion of the Atlantic Ocean, made of complex morphologies such as plateaus, seamounts, canyons, and rifts at water depths ranging from ~700 to 4000 m (Jovane *et al*., 2019). In the last decades it is becoming strategic for potential mineral exploitation of ferromanganese crusts (Montserrat *et al*., 2019; Benites *et al*. 2020; Bergo *et al*., 2021). However, the fact that deep sea is turning an official source of raw materials deserves an environmentally sustainable management (Guilhon *et al*., 2020) that may only be achieved with detailed biological diversity studies. The RGR is in the oligotrophic Subtropical South Atlantic Gyre (Perez *et al*., 2012), under a regenerated production regime *(sensu* Dugdale & Goering, 1967, Metzler *et al*., 1997), and minor contribution of diazotrophic microorganisms (Sohm *et al*., 2011). The main reservoir of new nutrients is trapped below the permanent thermocline, in the South Atlantic Central Water (SACW). The typical tropical profile (reference) that dominates the biogeochemical scenario in the whole area shows a deep chlorophyll maximum layer (DCM) as the result of photoadaptation of picoautotrophs to low-light conditions rather than biomass accumulation (Cullen, 2015). The aims of this study were to describe planktonic microbial communities over the RGR water column, specifically by (1) determining the picoplanktonic abundance, (2) assessing the diversity, structure, predict functions and composition of archaeal and bacterial communities, and (3) investigating oceanographic features setting and impact on the microbial community composition. To achieve this, we sampled seawater from several stations across the RGR, and combined flow cytometry and next-generation sequencing.

## MATERIALS AND METHODS

### Sampling Strategy

Seawater samples were collected at oceanographic stations along the RGR during the summer Marine E-Tech expedition (January - February 2018, more details in Jovane *et al*., 2019) onboard the R/V *Alpha Crucis* from the University of São Paulo (Figure 1). Sampling depths were defined according to the temperature (°C), salinity, oxygen (mg l^−1^), fluorescent dissolved organic matter (*f*DOM, QSU), chlorophyll and phycoerythrin fluorescence (RFU) profiles provided by a combined Sea-Bird CTD/Carrousel 911 5l Niskin rosette system, in order to sample the distinct water masses (TW - Tropical Water, SACW, AAIW - Antarctic Intermediate Water, and UCDW – Upper Circumpolar Deep Water).

**Figure 1.**
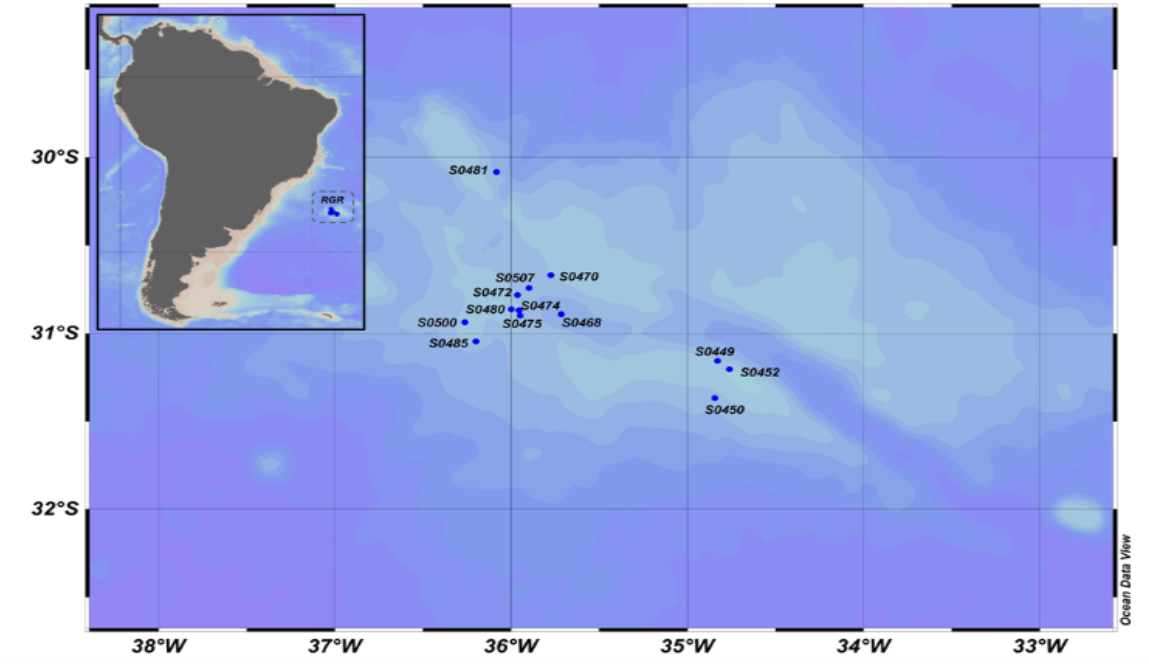
Sampling map on the Rio Grande Rise in the South Atlantic Ocean. Location of the RGR in relation to the Brazilian coast and oceanographic stations sampled in January - February 2018 (blue dots).

Seawater (9L) was filtered using a peristaltic pump through 0.22 μm membrane pores (Sterivex, Millipore, MA) for molecular biology. Triplicate samples (1.5 ml) for flow cytometry were stored into cryovials and immediately preserved with Sigma-Aldrich glutaraldehyde solution (0.1% of final concentration), and flash-frozen in liquid nitrogen for a few minutes to fix. All samples for biological analysis were stored at – 80°C for 30 days until analysis. Samples for inorganic nutrients (nitrate, nitrite, phosphate, and silicate) were obtained from the filtered 0.22-μm water, kept frozen (−20°C) and, on land, determined using a flow injection auto-analyzer (Auto-Analyzer 3, Seal *Inc*.)

Chlorophyll-a (chl-a) was also measured later in the lab by fluorometric techniques. For this between 1 – 5L of seawater sampled from the TW and DCM were vacuum filtered through 0.7 μm fiber glass filters (Whatman GF/F) and kept frozen in liquid nitrogen until laboratory analyzes. On land, the filters were extracted in 90% acetone and analyzed in Turner 10AU fluorimeter using the methods in Welschmeyer (1994). The chlorophyll fluorescence (RFU) from by the CTD was then converted into biomass concentration (μg L^−1^) applying a linear regression between the two measurements (r^2^=0.91, p<0.01, Supplementary Material).

### Flow Cytometry Analysis

Cell abundance of *Synechococcus spp., Prochlorococcus spp*., picoeukaryotes (≤ 2 μm), nanoeukaryotes (2-5 μm) and heterotrophic prokaryotes were determined according to Marie *et al*. (1999) using an Attune NxT flow cytometer (Thermo Fisher-Scientific) equipped with a syringe pump for quantitative volume sampling and two flat-top lasers: blue and red (488nm e 640nm respectively). Cells of *Prochlorococcus*, *Synechococcus*, picoeukaryotes and nanoeukaryotes were quantified based on autofluorescence in red (BL3 - intracellular concentration of chlorophyll, 4.560 nm) and orange (BL2 - intracellular concentration of phycoerythrin, 530 ± 30 nm) wavelengths simultaneously. To reduce the noise, a threshold of 500 was applied to BL3. After analyzing the autotrophs, SYBR Green I (Invitrogen Life Technologies, USA) was added, and the samples were dark incubated at room temperature for 15 min (Marie *et al*., 1999). Flow cytometry acquisition for heterotrophic bacteria was triggered on BL1 (530 ± 30 nm) with a threshold value of 500 and 90° side scatter. Data were analyzed using Flowjo software^®^ (https://www.flowjo.com/solutions/flowjo).

### DNA Extraction, 16S rRNA gene Amplification and Illumina Miseq Sequencing

DNA extraction was performed by using the DNeasy PowerWater (Qiagen, EUA) according to the manufacturer’s instructions. Negative (no sample) extraction controls were used to ensure the extraction quality. DNA integrity was determined after electrophoresis in 1% (v/v) agarose gel prepared with TAE 1X (Tris 0.04M, glacial acetic acid 1M, EDTA 50mM, pH 8), and staining with Sybr Green (Thermo Fisher Scientific, São Paulo, Brazil). DNA concentration was determined using the Qubit dsDNA HS assay kit (Thermo Fisher Scientific, São Paulo, Brazil), according to the manufacturer’s instructions.

Before sending samples for preparation of Illumina libraries and sequencing, the V3 and V4 region of the 16S rRNA gene was amplified with the primer set 515F (5’–GTGCCAGCMGCCGCGGTAA-3’) and 926R (5’–CCGYCAATTYMTTTRAGTTT-3’), (Parada *et al*., 2016) to check for the amplification of 16S using the extracted DNA. Negative (no sample) extraction controls were used for PCR amplification and the Illumina sequencing to check for the presence of possible environmental contamination (Sheik *et al*., 2018). Illumina DNA libraries and sequencing were performed at MR DNA (www.mrdnalab.com, Shallowater, TX, United States) on a MiSeq platform in a paired-end read run (2 × 250bp) following the manufacturer’s guidelines. Sequencing outputs were the raw sequence data.

### Flow Cytometry Data and Statistical Analysis

Sampling maps and graphs of the hydrographic conditions of the water column and picoplankton carbon distribution for north stations were generated using Ocean Data View^®^ with the DIVA gridding algorithm for variable resolution in a rectangular grid (Schlitzer, 2016). All data were used for statistical analyses with the R software (version 3.3.1).

Cell abundance was transformed (log(x+1)) and an nMDS (non-metric multidimensional scaling) using the Bray-Curtis similarity index was performed to look for patterns in the data. Environmental parameters were fitted to the nMDS using the envfit function of the vegan package with 999 permutations (Oksanen *et al*., 2016).

### Sequencing Data Analysis, Statistical Analysis and Data Visualization

Processing of 16S rRNA gene sequences followed a QIIME (Quantitative Insights Into Microbial Ecology) and Uparse pipeline (Edgar, 2013). The program PEAR was first used to merge the multiplexed, paired-end sequences and the program Usearch version 8.1.1861 was used to filter the merged sequences with a minimum error of 1.0. Chimeric sequences were removed, and the Operational Taxonomic Units (OTUs) were clustered based on 97% sequence similarity using Usearch. The reference Silva132 database was used to assign the taxonomic classification of the OTUs.

Statistical analyses were conducted using R software with the specific packages ggplot2 and vegan (Venables & Smith, 2003). Alpha-diversity indexes (number of OTUs, Shannon and Chao1) were calculated using the QIIME script alpha_diversity.py, and differences in alpha-diversity estimates between groups of samples were tested using the Student’s t-test. The beta diversity, based on the UniFrac weighted phylogenetic index, was calculated using the QIIME script core_diversity_analyses.py (Lozupone & Knight, 2005), visualized via non-metric multidimensional scaling (nMDS). Differences in the communities between the epi- (0-200 m), meso- (200 – 1000 m) and bathypelagic (1000 – 4000 m) zones were tested by Permutational Multivariate Analysis of Variance (PERMANOVA) using the function adonis from the package vegan, based on the UniFrac distance with 999 permutations.

The Indicator Species analysis was performed to identify significant taxonomic indicators (Cáceres *et al*., 2010) within the distinct water masses (TW - Tropical Water, SACW - South Atlantic Central Water, AIW - Antarctic intermediate Water, and CSW - Circumpolar Superior Water) from RGR. The analysis was conducted on OTU counts excluding OTUs < 20 reads. The results were visualized as networks by means of the igraph R package (Csárdi & Nepusz, 2006). Predicted microbial functional groups were identified using the Functional Annotation of Prokaryotic Taxa (FAPROTAX) 1.2.1 database (Louca *et al*., 2016). Statistical tests were considered significant at p < 0.05. FAPROTAX extrapolated functions of cultured prokaryotes (identified at the genus or species level) to the rest of the prokaryotic genus to estimate putative functions. Raw sequence data generated for this are publicly available in the National Centre for Biotechnology Information (NCBI) database under the BioProject PRJNA714894.

## RESULTS

### Water Column and Environmental Factors

Seawater temperature ranged from 2 - 25°C, with the coolest temperatures near the bottom of the ocean at station 481, and the warmest temperatures in shallow waters at station 480 (Supplementary Figure 1). Salinities ranged from 34.26 PSU at station 481 to 36 PSU at station 450 (Supplementary Figure 1). The temperature-salinity signatures indicate the presence of four main pelagic water masses: Tropical Water - TW (T>20°C and S>36 PSU) at ca. 150 m depth, South Atlantic Central Water - SACW (8.7°C<T<20°C and 34.6 PSU <S<36 PSU) between 150 m and 500 m depth, Arctic Intermediate Water - AIW (3.4°C<T<8.7°C and 34.3 PSU <S<34.6 PSU) at 500 m and 1500 m depth, and Circumpolar Deep Water - CDW (T?3°C and 34.5 PSU <S<34.8 PSU) at 1300 m and 1600 m depth.

Relatively lower temperatures and higher dissolved oxygen (>7 mg/L) were associated with fluorescence peaks (>1.5 RFU), for example, at station 480 the 18°C isotherm was found 115m deep, resulting in a 2.64 RFU peak fluorescence. Chlorophyll fluorescence ranged from 2.73 RFU, at station 480, to 0.22 RFU at station 449, with the DCM layer occurring in the depth of 80 to 150 m (Supplementary Table 1 and Supplementary Figure 1).

The nutricline was also well-defined and coincident with or immediately adjacent to the SACW upper limit at samples, except for nitrite (Supplementary Figure 2). The concentrations of nitrate and phosphate and silicate ranged between undetected concentrations of both nutrients up to a maximum of 42.94 μM and 3.1 μM, respectively. High CDOM (>0.1 mg L^−1^) coincided with the presence of fluorescence peaks at all stations. The CDOM varied between 0.02 to 0.19 mg L^−1^ (Supplementary Table 1).

Stations representing the different pelagic zones and water masses were clearly distributed based on their environmental and physical characteristics in our PCA analysis (Supplementary Figure 3). Three PCA components explained 83.03% of the sample variability based on environmental parameters (temperature, salinity, oxygen, chlorophyll, nitrite, nitrate phosphate, silicate and CDOM). The first component (PC1) was negatively correlated with temperature and salinity and positively correlated with nitrate, which may reflect the influence of the cold and nutrient rich SACW. The second axis (PC2) was negatively correlated with chlorophyll, while the third component (PC3) showed a high negative correlation with nitrite and positively correlated with oxygen (Supplementary Table 2).

### Pico- and Nanoplankton abundance

*Prochlorococcus* were dominant in the bottom of the TW and in the upper limit of SACW, ~100 m depth. The highest abundances were recorded at stations 485, 481 and 480, with 5.12 x 10^4^, 4.82 x 10^4^ and 4.3 x 10^4^ cell mL^−1^, respectively. Generally, in surface waters, few cells were recorded, however, *Prochlorococcus* corresponded to 72.4 %o of the total autotrophic cells (Supplementary Table 3 and Supplementary Table 4).

*Synechococcus* were more abundant in the surface layers, especially in the TW (Supplementary Table 3). Maximum cell numbers were counted above DCM at all stations, and the highest abundances were recorded at 11 and 5 m depth in the stations 475 and 507 (7.22 x 10^3^ cell.mL^−1^ and 4.77 x 10^3^, respectively). Decreasing abundances were observed with depth and *Synechococcus* corresponded to 18 % of the total autotrophic cells (Supplementary Table 3 and Supplementary Table 4).

A similar vertical and spatial distribution was observed for photosynthetic picoeukaryotes and nanoeukaryotes. They were more abundant in the depth between 90 and 120 m in all stations. The maximum concentrations (4.68-4.06 x 10^3^ cell.mL^−1^, station 481) were found at 100 m depth, whereas lower abundance (<5.10 x 10^1^ cell.mL^−1^) were recorded below depths of 450 m. Picoeukaryotic populations corresponded to 9.4 % of the total autotrophic cells (Supplementary Table 3 and Supplementary Table 4). Nanoeukaryotes were more abundant at stations 450 (9.3 x 102 cell.mL-1), 468 (8.61 x 102 cell.mL-1) and 449 (8.43 x 102 cell.mL-1). Higher cell concentrations were recorded at ca. 100 m. Nanoeukaryotes was (1.5 %) the less abundant group of the total autotrophic cells (Supplementary Table 3 and Supplementary Table 4).

Heterotrophic bacteria dominated (averaging 99%) the microbial community (Supplementary Table 4). The higher abundances >10^5^ cell mL^−1^ occurred at stations 449 and 450. The highest concentration was in the TW, at stations 449 (8.52 x 10^8^ cell.mL^−1^, 120 m) and 450 (8.42 x 10^8^ cell.mL^−1^, 43 m). Other samples had concentrations four orders of magnitude lower (Supplementary Table 3 and Supplementary Table 4).

The spatial and vertical distribution of total picophytoplankton and heterotrophic bacteria were similar, with higher abundance of both groups at surface tropical oligotrophic waters (Figure 2F1 and F2). The higher picophytoplankton abundances were at stations 481, 485, 468 and 475, corresponding with the transition between TW and SACW.

**Figure 2.**
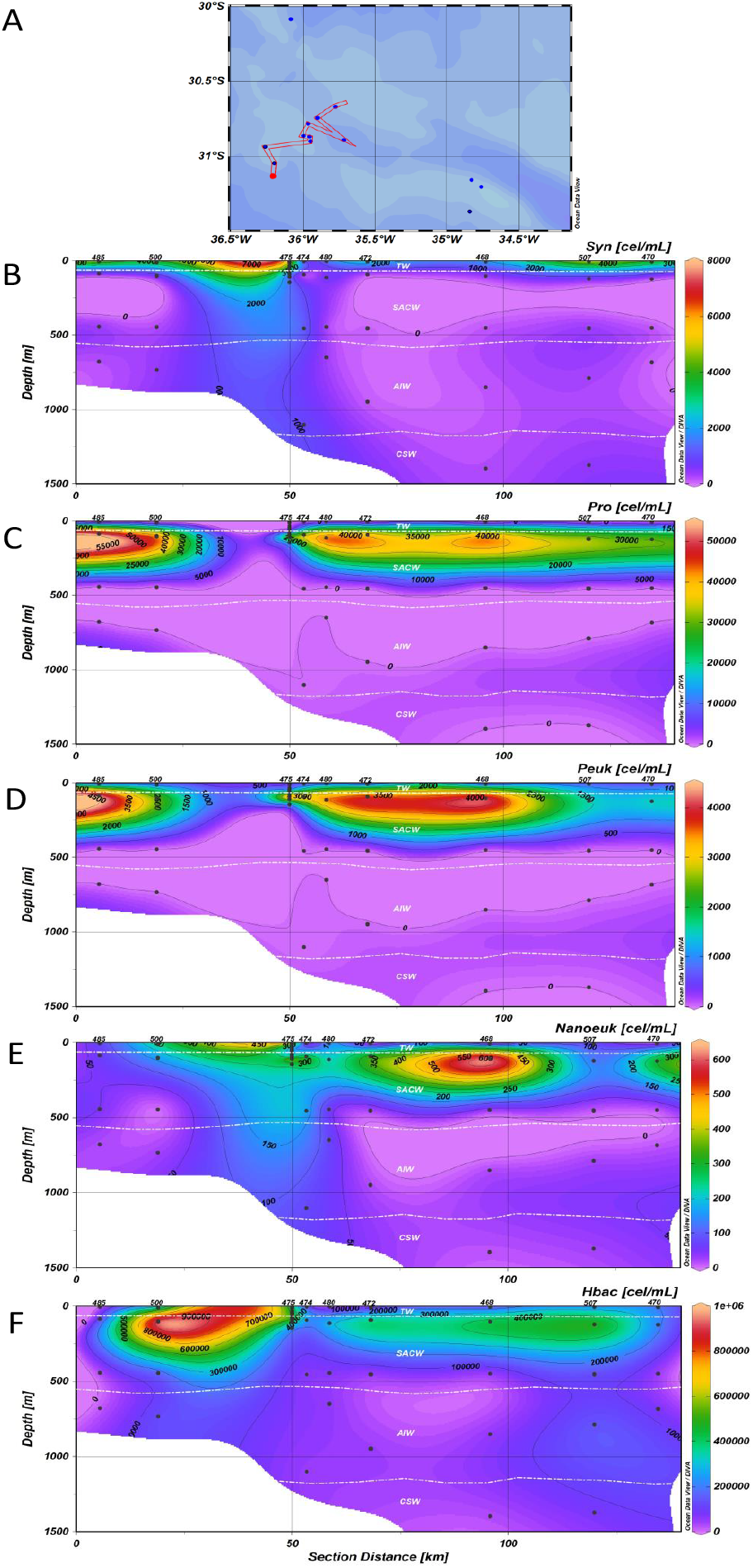
Picoplankton abundance along the main sampling area in the Rio Grande Rise: (A) West stations, depth profiles of cells (cell mL^1^) distribution of (B) *Synechococcus* (Syn), (C) *Prochlorococcus* (Pro), (D) Picoeukaryotes (Peuk), (E) Heterotrophic bacteria (Hbac), and (F) Total Picophytoplankton. Additional stations to the east were also used to statistics.

Analyses using nMDS showed the picoplankton distribution over the pelagic zones and across environmental parameters, including temperature, salinity, fluorescence, and inorganic nutrients, as well as the water masses (Figure 3). The biological groups correlated significantly with depth, temperature, salinity, oxygen and fluorescence (p < 0.005). The plankton community distribution was explained by the pelagic zone according to the PERMANOVA test (r^2^ = 17%, p = 0.003).

**Figure 3.**
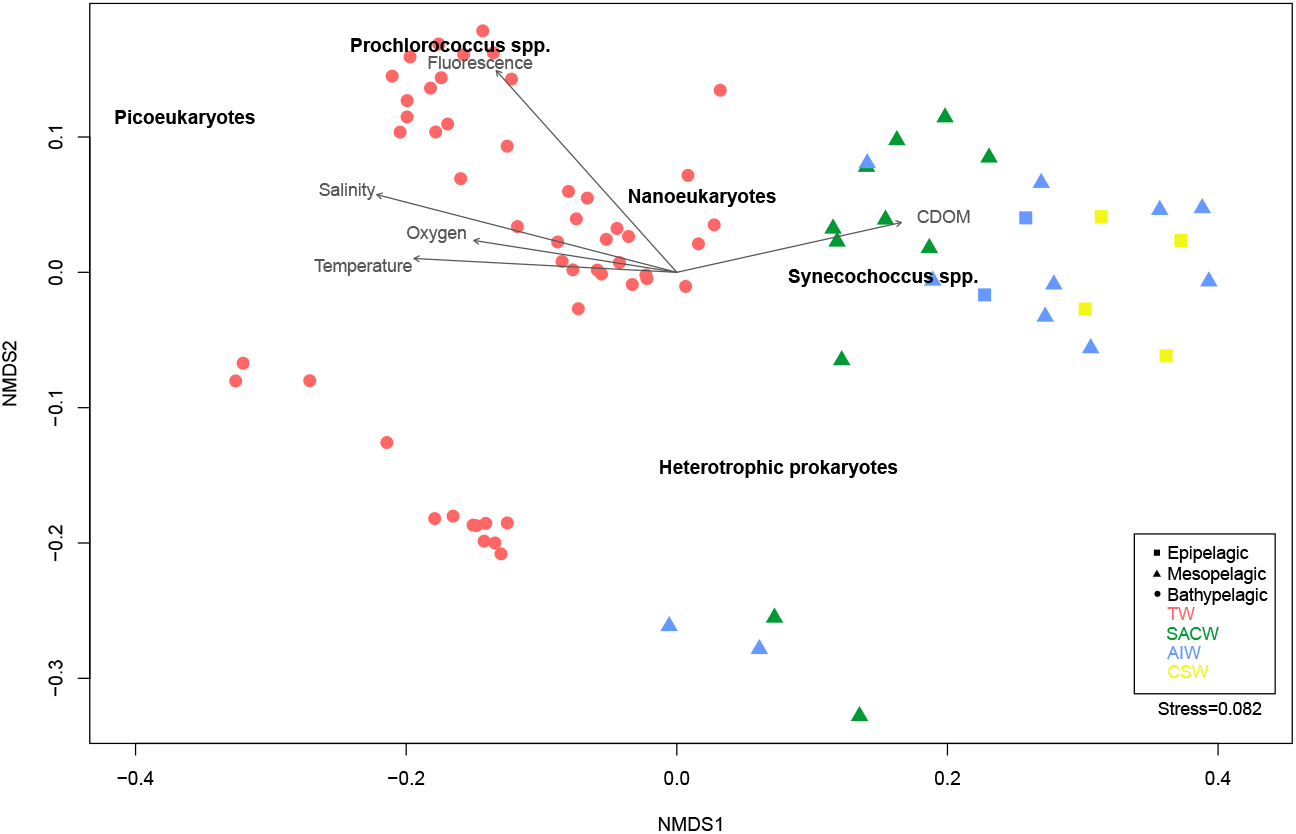
Non-metric multidimensional scaling (nMDS) ordination of the Picoplankton abundance from the RGR (stress = 0.082, r^2^ = 0.996). Circles represent samples collected in the epipelagic zone, triangles represent samples collected in the mesopelagic zone, and squares represent samples collected in the bathypelagic zone. Different colors represent the water masses in the RGR: Red is Tropical Water (TW), green is South Atlantic Central Water (SACW), blue is Antarctic Intermediate Water (AIW) and yellow is Circumpolar Superior Water (CSW). Each arrow shows one environmental gradient significantly correlated to the ordination (envfit, p < 0.005). The arrows point to the direction of the most rapid change in the environment (direction of the gradient) and its length is proportional to the correlation between ordination and environmental variable (strength of the gradient).

### Microbial Alpha and Beta Diversity Estimates

A total of 2,227,889 sequences was retrieved from 32 samples and ranged from 23,650 to 100,897 per sample. After removing chimeras, sequences were clustered into 2,431 operational taxonomic units (OTUs) (0.03 cut-off). After filtering other contaminant groups, a total of 2,329 OTUs were obtained. Alpha diversity indices were not statistically different among the pelagic zones (Kruskal-Wallis one-way ANOVA, p>0.005) (Supplementary Figure 4).

Beta diversity was explored by weighted Unifrac distances and ordered by nMDS. The nMDS shows that microbial communities are clearly separated by epipelagic and, meso- and bathypelagic zones (stress:0.02; PERMANOVA, r^2^= 0.28, p=0.001), and post-hoc analysis revealed significant differences between each possible pairwise comparison between pelagic zones, except for meso- and bathypelagic zones (Figure 4 and Supplementary Table 5). Permuted analyses of beta-dispersion showed significant differences in the distance to the centroid (spread) among pelagic zones (betadisper, F= 3.42, p=0.004), indicating that differences in microbial communities could be due to differences in within-group dispersions instead of variation in centroid position.

**Figure 4.**
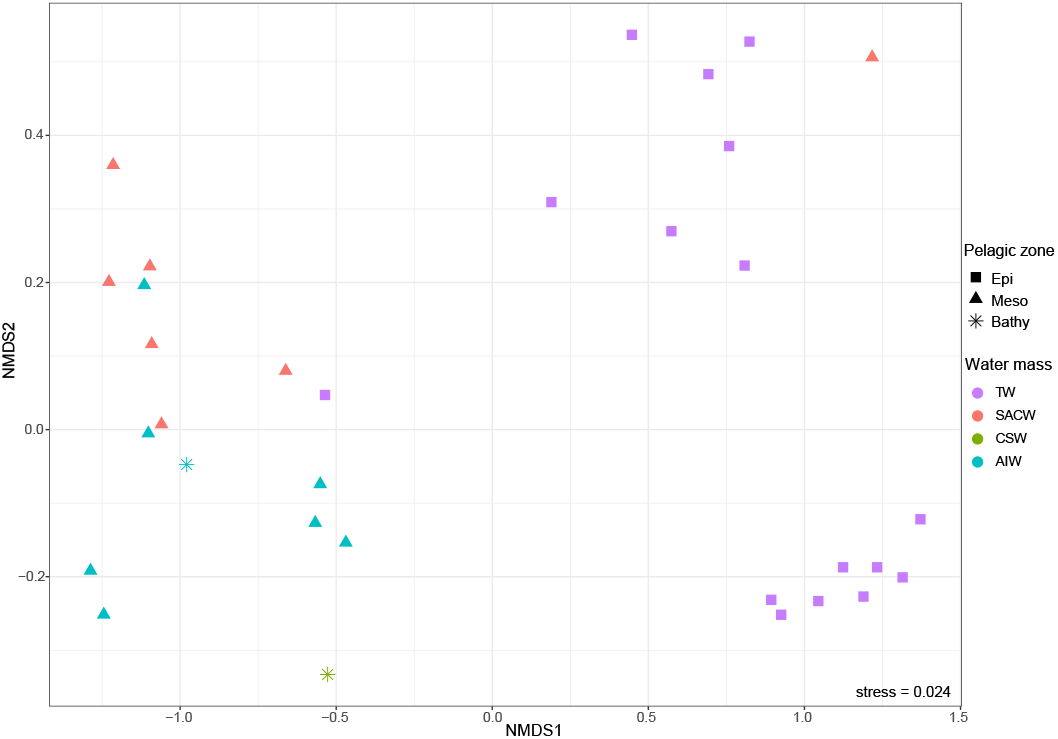
Non-metric multidimensional scaling (nMDS) ordination of the microbial community from pelagic zones in the RGR (stress = 0.024). Circles represent samples collected in the epipelagic zone, squares represent samples collected in the mesopelagic zone, and triangles represent samples collected in the bathypelagic zone. Different colors represent the waters masses: Red is Tropical Water (TW), blue is South Atlantic Central Water (SACW), green is Antarctic Intermediate Water (AIW), and yellow is Circumpolar Superior Water (CSW).

### Microbial Community Structure Across Oceanographic Features

To identify key oceanographic features in shaping microbial community composition, Spearman correlations were calculated, and only significant (p < 0.05) and strong correlations (r > - 0.5 or 0.5) were considered. In general, bacterial (OTU8, OTU25 and OTU156) and Archaeal (OTU4 and OUT11) OTUs prevalent in the epipelagic zone showed a positive correlation with fluorescence and nitrite and negative correlation with phosphate, silicate and depth. In contrast, bacterial (OTU14, OTU5, OTU7, OTU101, OTU10 and OTU644) and Archaeal (OUT44) OTUs prevalent in meso- and bathypelagic zones revealed negative correlations with fluorescence and nitrite and positive correlations with phosphate, silicate and depth. Dominant OTUs (OTU2, OUT98 and OUT156) in all pelagic zones revealed positive correlations with fluorescence and nitrite (Figure 5).

**Figure 5.**
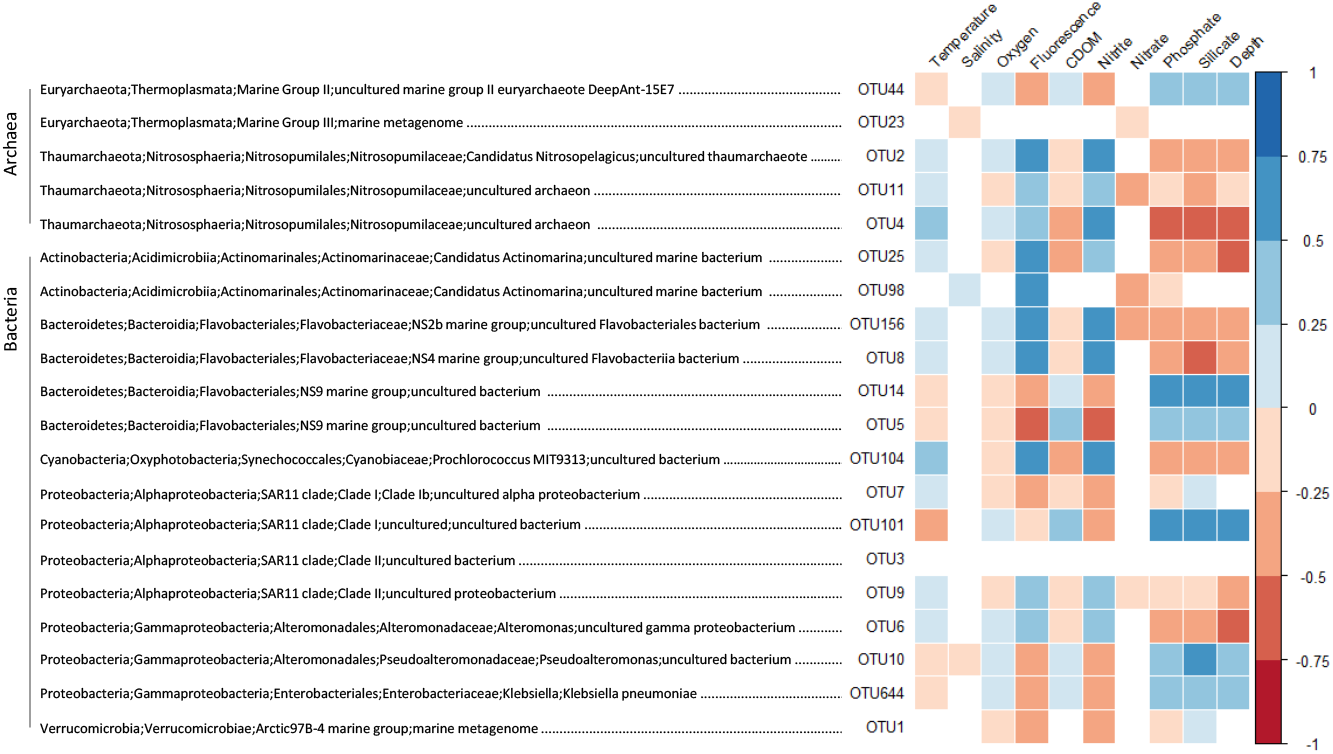
Spearman correlation between archaeal and bacterial abundant OTUs (>1% of relative abundance), and Temperature, salinity, oxygen, chlorophyll, density, turbidity, CDOM and phycoerythrin. Only parameters that exhibited p < 0.05 in correlations analysis are represented.

### Microbial community composition

The analysis of all sequences of the 16S rRNA gene showed that the communities were dominated by Bacteria (80%) rather than by Archaea (20%). At the phylum level, bacteria were dominated by Proteobacteria (69%, classes Gamma - and Alphaproteobacteria), Bacteroidetes (5.7%) and Cyanobacteria (3.8%) phyla. All archaeal communities were dominated by Thaumarchaeota (*Nitrosopumilales*), followed by Euryarchaeota (*Marine Group*) (Figure 6 and Supplementary Figure 5).

**Figure 6.**
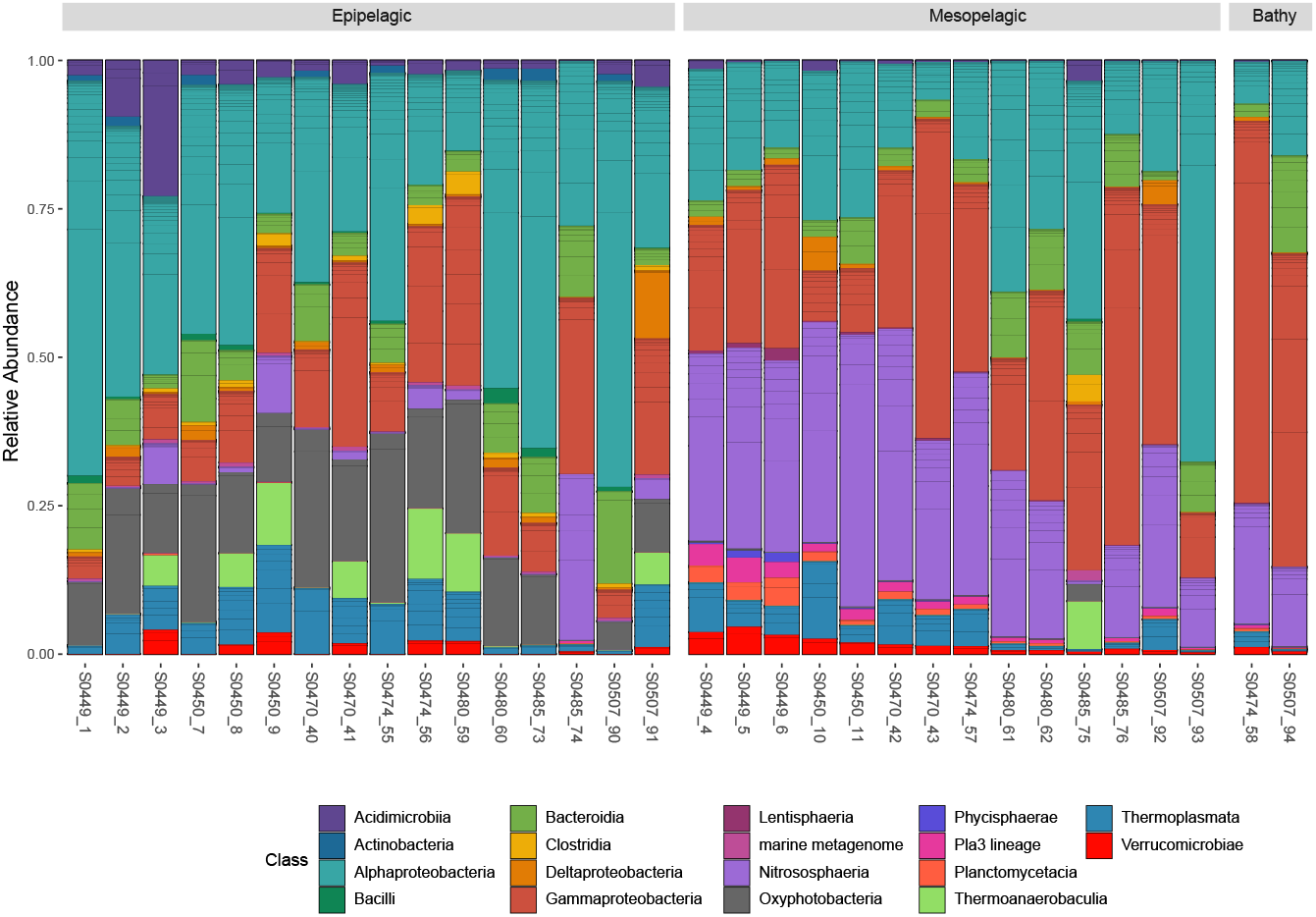
Relative abundance of bacterial and archaeal taxonomic groups at the class level on the Rio Grande Rise. Only groups with more than 0.1% abundance are represented. Depths and sites of each sample are represented. Sequences were clustered at 97% similarity and taxonomy was assigned by performing BLAST searches against the Silva database v. 132 (*E*-value ≤ 1*e*–5).

Most taxonomic groups varied in abundance or occurrence between the pelagic zones. For example, an increasing pattern in the relative abundance of *Alteromonadales* (Gammaproteobacteria class) and *Nitrosopumilales* (Nitrososphaeria class) with depth was observed. While *Synechococcales* were mainly found near the surface and DCM (Figure 6 and Supplementary Figure 5).

At the class level, the epipelagic zone (0-200m) was dominated by Alphaproteobacteria (66-13%, order SAR11 clade I), Gammaproteobacteria (30-5%, order Alteromonadales), Oxyphotobacteria (25-1%, order Synechococcales), Bacteroidia (16-4%, order Flavobacteriales), Thermoplasmata (15-1%, order Marine Group II) and Nitrososphaeria (28-1%, order Nitrosopumilales) (Figure 6). In the mesopelagic zone, the prevalent classes were Nitrososphaeria (45-1%, order Nitrosopumilales), Gammaproteobacteria (58-8%, order Alteromonadales), Alphaproteobacteria (66-6%, order SAR11 clade I), Bacteroidia (12-1%, order Flavobacteriales) and Thermoplasmata (12-1%, order Marine Group II) (Figure 6). The two samples from the bathypelagic zone (1000m-4000m) were dominated by OTUs affiliated to the Gammaproteobacteria (63-53%, order Alteromonadales), Nitrososphaeria (20-14%, order Nitrosopumilales), Alphaproteobacteria (15-6%, order SAR11 clade I) and Bacteroidia (14-2%, order Flavobacteriales) (Figure 6).

Microbial community composition among different water masses was further investigated by means of the Indicator Species analysis (Figure 7). We compared the relative abundance and relative frequency of each OTU to identify those specifically associated with only one water mass (unique) and those whose niche breadth encompasses all water masses (shared). The water masses CSW and TW harbored the highest sets of unique OTUs (n = 106 and n = 29, respectively). The unique OTUs for TW belonged to the SAR11 Clade I (n = 5), Clade II (n = 1), Clade IV (n = 1) and Flavobacteriales (n = 4). CSW unique OTUs mainly belonged to the Flavobacteriales and Cellvibrionales, as well as one belonging to Bacillales (OTU31 - class Bacilli), for which the functional prediction was manganese oxidation. A different pattern emerged for SACW and AIW, which harbored fewer unique OTUs (only n = 7 and n = 3, respectively), that belonged to Rhodobacterales and uncultured archaea (n = 1 and n = 2, respectively) for SACW and Planctomycetales, Francisellales and uncultured euryarchaeote for AIW. The SACW, CSW and AIW shared 3 OTUs that were related to the Nitrosopumilales (n = 2) and SAR11 clade II. The CSW shared 4 OTUs with TW (mostly Bdellovibrionales) and 1 OTU with AIW (Phycisphaerales). SACW shared 2 OTUs with AIW (both Nitrosopumilales) (Figure 7).

**Figure 7.**
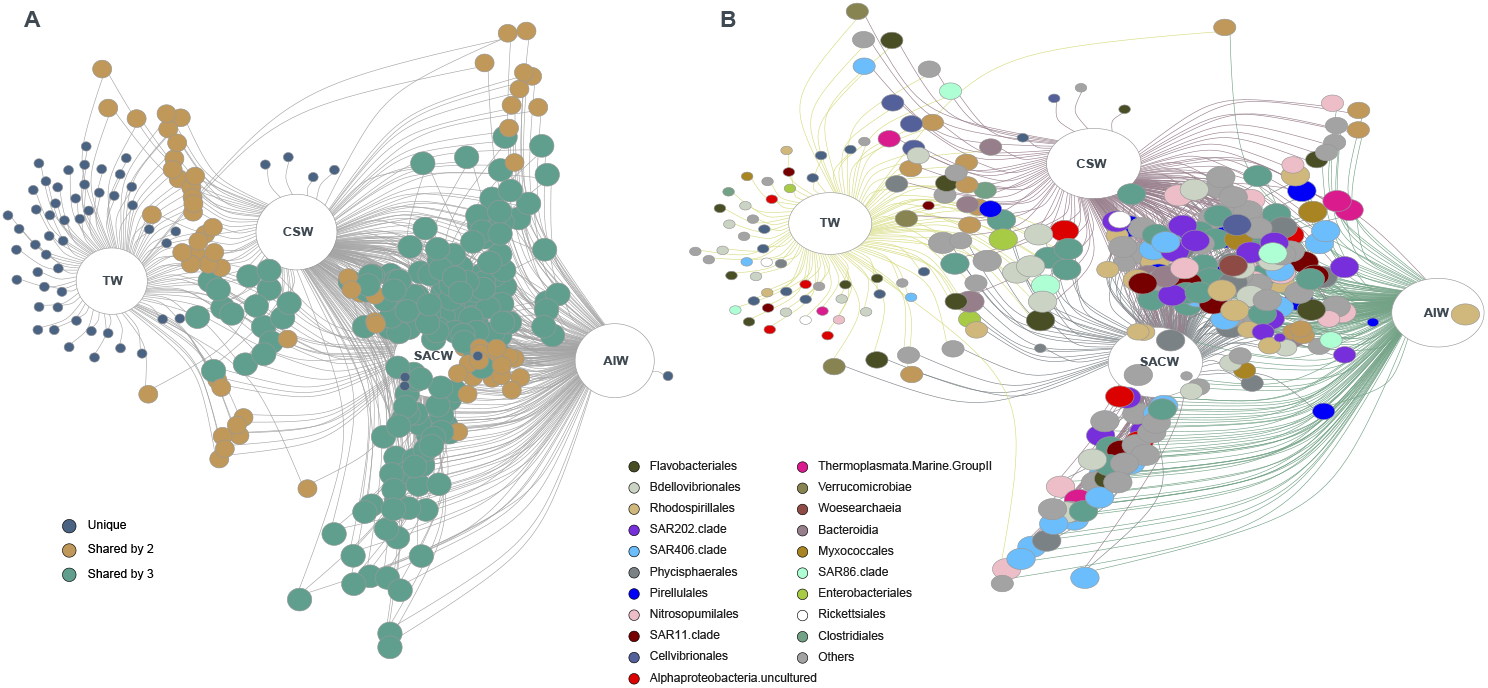
Results of the Indicator Species analysis for water masses from RGR. The network (A) indicates the correlation (represented by edges) of unique and shared OTUs (between 2, 3 and 4 different water masses presented as TW - Tropical Water, SACW - South Atlantic Central Water, AIW - Antarctic Intermediate Water and CSW-Circumpolar Superior Water). The network (B) highlights the taxonomic classification of the nodes that belonged to the most represented orders (i.e., those representing > 1% of relative abundance in at least one sample). Non-dominant taxa (i.e., < 1%) are reported as “Others”. The size of the nodes reflects their “fidelity” to the substrates, so that a large node indicates those OTUs always present in all the samples of that water mass.

### Predicted function

We assigned 806 out of 2,329 microbial OTUs (33.15%) to at least one microbial functional group using the FAPROTAX database (Supplementary Table 7). Nitrogen fixation, fermentation, aerobic chemoheterotrophy, methylotrophy, nitrification, aerobic ammonia oxidation, dark sulfur, sulfite and sulfide oxidation, dark thiosulfate oxidation, aromatic compound degradation, nitrate respiration and reduction, and anoxygenic and oxidizing photoautotrophy were functions predicted in all pelagic zones with differences in relative abundance between samples (Supplementary Figure 6). Main functions associated with microbial communities from the epipelagic zone were photoautotrophy, nitrite and nitrate denitrification, and manganese and iron respiration. The predicted functions in samples from meso- and bathypelagic zones were manganese, sulfur and sulfide oxidation, methanogenesis, methanotrophy, nitrite and nitrate ammonification, and degradation of carbon compounds (Supplementary Figure 6).

## DISCUSSION

### Distribution patterns of picoplankton and its link with physicochemical parameters

Marine microorganisms have singular biogeographic distribution patterns in the ocean, particularly under morphological features such as seamounts and rises that affect spatial distributions and concentration of the picoplankton community (Milici *et al*., 2016; Giljan *et al*., 2020; Rocke *et al*., 2020).

Picoeukaryotes dominated the (> 77 %) the picophytoplankton across the RGR, as usually reported in mesopelagic ecosystem (Agustí *et al*., 2015; Giner *et al*., 2020). They dominated in and below the DCM, mainly associated with the SACW, with higher concentrations of nitrate and phosphate. Pajares *et al*., (2019) suggested that these organisms prefer nutrient-rich waters. The occasional detection of metabolically active and significant abundances of picoeukaryote groups in the dark ocean has been previously described (Agustí *et al*., 2015; Giner *et al*., 2020). This could be due to their attachment to rapidly sinking particles (Agustí et al, 2015) or migration in the water column in response to different gradients of light and nutrients to improve their growth conditions (Raven, 1998). Despite this, some picoeukaryotes may be mixotrophs (Faure et al, 2019). In this study, picoeukaryotes at and below the nutricline can be related to the resident populations that use nitrate from deeper waters (Bergo *et al*., 2017).

The abundance of *Prochlorococcus* was higher at ~100 m depth, close to the nutricline, which might indicate an ecotype adapted to low light intensity (Garcia-Robledo *et al*., 2017). In contrast, they were less abundant (sometimes even absent) in surface (<50m) and deeper (>450m) layers compared to other picophytoplankton groups. According to Lange *et al*. (2018), *Prochlorococcus* is expected to have the highest abundance within the picoautotrophs in oligotrophic waters, and yet we did not confirm this assumption. Previous studies have described that low concentrations of photosynthetic pigments with the *Prochlorococcus* ecotypes in surface waters may lead to an underestimation of cell concentrations by the cytometer (Partensky *et al*., 1999; Gérikas-Ribeiro *et al*., 2016a). Nevertheless, the observed mean cell concentrations were in the same order of magnitude as previous studies (Guo *et al*., 2014; Bergo *et al*., 2017), and this underestimation does not affect the spatial distribution patterns of picoplankton reported in this study. The abundance of *Synechococcus* was higher in the epipelagic zone, above the DCM associated with the relative warm and nutrient poor TW. Normally, the abundance of these cyanobacteria is lower in deep oceanic oligotrophic waters decreasing with depth (Wang *et al*., 2011). Colder and nutrient-rich waters (including trace metals) inhibit *Synechococcus* growth (Wang *et al*., 2011).

Heterotrophic bacteria dominated the microbial community in the water column, with one order of magnitude more abundant than the other pico-sized groups. Cotner & Wetzel (1992) suggest that heterotrophic bacteria dominate in oligotrophic waters in comparison to picophytoplankton. The related distribution between heterotrophic bacteria and picophytoplankton abundance in the water column reveals connection among the picoplankton communities, the production of organic matter and nutrient cycling in the RGR. This was expected particularly in the euphotic zone considering that the growth of heterotrophic bacteria is partially supported by dissolved organic compounds produced by phytoplankton cells, and the remineralization of inorganic nutrients by heterotrophic bacteria stimulates the phytoplankton growth (Linacre *et al*., 2015). Moreover, other sources of dissolved organic compounds may sustain the heterotrophic communities, for instance, viral lysis, seasonal blooms and senescent cells (Buchan *et al*., 2014) which explains their dominance in deeper aphotic layers.

In general, the density of autotrophic and heterotrophic picoplankton populations was comparable to previous studies in the South Atlantic Ocean (Bergo *et al*., 2017; Gérikas-Ribeiro *et al*., 2016a; Gérikas-Ribeiro *et al*., 2016b). Excluding the heterotrophs, the *Prochlorococcus* dominated the autotrophic groups in our study (72.38%), followed by *Synechococcus* (16.77%), picoeukaryotes and picoeukaryotes (9.39%). Besides photoadaptation by increasing the intracellular photosynthetic pigments, *Prochlorococcus* benefit from the small flux of nitrate (Berube *et al*., 2015) across the main pycnocline which explains their dominance among the picophytoplankton.

Although the abundance of the picoplankton varied vertically across sampling depths perhaps due to species-specific differences in photoadaptation and nutrient uptake efficiency, it also varied among stations due to submesoscale (1-5km) changes of turbulence and environmental conditions. The differences of microbial abundance and diversity among pelagic zones and water masses here observed, contribute for niche diversity and their spatial distribution.

### Microbial Community Composition and Diversity

Studies have shown that picoplankton microbial communities are structured by water masses (Agogué *et al*., 2011; Celussi *et al*., 2018), pelagic zones (Walsh *et al*., 2016) and depth (Mestre *et al*., 2017). Deep-sea water masses were suggested as “bio-oceanographic islands” for prokaryotic plankton (Agogué *et al*., 2011), and that dark ocean microbiome is primed by the formation and the horizontal transport of water masses (Frank *et al*., 2016). Besides that, submarine mountains or elevations that divide the deep ocean into deep-oceanic basins may act as “ecological barriers” for prokaryotic communities (Salazar *et al*., 2016). Indeed, our NMDS analysis showed that all samples were highly divided into pelagic zones and water masses with a strong correlation between the microbial community and depth, nitrite, phosphate, and silicate. The different nutrient availability in each pelagic zone and water masses may be an important oceanographic feature in shaping the microbial communities in the RGR. However, our results showed there are no differences in bacterial and archaeal alpha diversity among the pelagic zones and water masses.

Microbial assemblages in the water column from the RGR showed the dominance of the classes Alphaproteobacteria and Gammaproteobacteria, within the Proteobacteria phylum, as described for other seamounts (Kato *et al*., 2018; Giljan *et al*., 2020) and the South Atlantic Ocean (Coutinho *et al*., 2020). The dominant bacterioplankton across the RGR consisted of phototrophic (*Synechococcales*) and heterotrophic (SAR11 Clade) members, which are both well-established marine oligotrophs regions with streamlined genomes and small cell size (Partensky *et al*. 1999; Giovannoni *et al*. 2017). *Synechococcales* are the dominant photosynthetic group in many epipelagic regions (Coutinho *et al*., 2020), whereas the SAR11 clade is the most abundant heterotrophic group in mesopelagic environments (Frank *et al*., 2016) and in different water masses (Celussi *et al*., 2018). The dominance of SAR11 in the whole water column from RGR could be due their capacity to respond to carbon limitation by metabolizing endogenous carbon (e.g. methane and methanol) in the dark and using proteorhodopsin to provide proton motive force in the light (Giovannoni *et al*., 2017). In our study, SAR11 Clade I represented the largest part of unique oligotypes in TW and SAR11 Clade II were shared between SACW, CSW and AIW.

Aerobic heterotrophic groups adapted to high-light environments, such as Flavobacteriales and Verrucomicrobiales, were mostly found in the euphotic zone over the RGR. Previous studies have shown that flavobacterial phylotypes are one of the dominant groups responding to phytoplankton blooms (Teeling *et al*., 2012, 2016). This response consists of a succession of particular clades, namely NS9, NS4, and others, related to the consumption of algal-derived organic matter, and gene repertories that allow the uptake and use of specific biopolymers during and after the bloom (Teeling *et al*., 2016; Unfried *et al*., 2018).

The relative abundance of *Gammaproteobacteria* in the RGR increased with depth as reported previously for the North (Lauro & Bartlett, 2008) and South Atlantic Ocean (Coutinho *et al*., 2020). Within *Gammaproteobacteri*a, members of the order *Alteromonadales* (mostly represented by Pseudoalteromonas and Alteromonas) were abundant in meso- and bathypelagic waters in this study. These marine heterotrophs play important roles in the biodegradation of marine organic matter, such as organic nitrogen and phosphorus mineralization (Li *et al*., 2015). In our dataset, the relative abundance of Alteromonadales from bathypelagic water was significantly and negatively corelated with nitrite (r = −0.55, p < 0.01), suggesting that the availability of nitrogen compounds may be a key determinant for these bacteria in the deep-sea. Besides that, members of this family have been previously reported in Fe-Mn deposits and suggested to contributed to manganese (Wu *et al*., 2013) and iron oxidation (Li *et al*., 2020).

Archaea is known to be an important component of the prokaryotic communities in meso- and bathypelagic water masses in the oceans (Grzymski *et al*., 2012) and in the Fe-Mn deposits (Shulse *et al*., 2017; Kato *et al*., 2018). Previous studies have described that dissolved ammonia in seawater is an important nutrient in the Fe-Mn deposit regions (Shulse *et al*., 2017; Kato *et al*., 2018). Based on the recovered OTUs and the functional predictions performed in this study, a high proportion of chemolithoautotrophic ammonia-oxidizing Archaea (Nitrososphaeria class) was detected in deep waters from RGR, especially in the mesopelagic waters. Nitrososphaeria represented the largest part of unique oligotypes shared between SACW, CSW and AIW in the RGR. The large proportion of these archaea in deep waters may reflect the importance of reduced substrates such as ammonia in the dark ocean (Shulse *et al*., 2017). Marine Group II (MG II) represents the most abundant archaeal group in ocean surface waters (Zhang *et al*., 2015) but in this study, it was prevalent in meso- and bathypelagic waters. It has been suggested that members of deep-sea MG II have unique patterns of organic carbon degradation with a chemoheterotrophic lifestyle (Rinke *et al*, 2019).

Recently, Bergo *et al*. (2021) suggested that carbon and nitrogen microbial metabolisms are likely important for deep-sea Fe–Mn crusts from the RGR. Despite the FAPROTAX limitations of bacterial and archaeal functions prediction, the main metabolisms predicted in the pelagic zones from the RGR were related to carbon, nitrogen and sulfur cycles. In addition, the future mining/dredging of Fe–Mn crusts may change the physical structure of the seamount/rise and would potentially impact water circulation in the deep-sea (Orcutt *et al*., 2020). These disturbances can consequently affect microbial ecology and functions, and biogeochemical cycles (Niner *et al*., 2018, Orcutt *et al*., 2020). However, little is known about water circulation at RGR (Montserrat *et al*., 2019), so it is difficult to know how this alteration could affect the area.

In conclusion, the overlap of water masses in the pelagic zone creates different niches that increase the diversity of microbial communities and determine their spatial distribution and ecological function. Intermediate temperatures in the mesopelagic zone, with SACW and AIW, creates a strong environmental driver for community structure and diversity, with a higher presence of chemosynthetic archaea than heterotrophic bacteria in oligotrophic waters of the RGR. The predicted functions of the deep-sea microbiome indicate that populations associated with nitrogen, carbon and sulfur are likely important contributors to the ecological process occurring in the RGR. However, further investigation is necessary to understand to which extent the pelagic microbial community affects the redox potential of deeper layers and ultimately the development of metallic crusts in the RGR in the South Atlantic Ocean.

## Funding

This study was funded by the São Paulo Research Foundation (FAPESP), Grant number: 14/50820-7, Project Marine ferromanganese deposits: a major resource of E-Tech elements, which is an international collaboration between Natural Environment Research Council (NERC, UK) and FAPESP (BRA). JCN received a scientific initiation fellowship from Programa Institucional de Bolsas de Iniciação Científica — CNPq/ PIBIC (Process number 156954/2018-4). NMB was financed by a Doctoral fellowship from the *Coordenação de Aperfeiçoamento de Pessoal de Nível Superior* - Brasil (CAPES) - Finance Code 001. VHP and LJ are supported by FAPESP gran number 18/17061-6.

## Conflict of Interest

The authors declare no conflict of interest.

## Availability of data and material

The sequencing reads generated for this study can be found in the National Centre for Biotechnology Information (NCBI) database under BioProject PRJNA714894.

## Acknowledgments

We are grateful to the captain and the crew of the Research Vessel *Alpha Crucis*, and scientists who joined the expedition Marine E-Tech RGR1 for their cooperation in sample collection. We would like to thank Linda Waters for the English language review. Sincerest gratitude goes out to Carolina L. Viscarra and MSc. Mariana Benites for their scientific support on board. Concentrations of inorganic nutrients incorporated in this study were provided through the hard work of M.Sc Mayza Pompeu and Giulia Campos. A huge thank you to LECOM’s research team and M.Sc Rosa C. Gamba for their scientific support.

## Author contributions

Juliana C. N. Ferreira: investigation, methodology, formal analysis, visualization, writing - original draft, writing - review & editing. Natascha M. Bergo: investigation, methodology, formal analysis, visualization, writing - original draft, writing - review & editing. Pedro M. Tura: investigation, writing - review & editing. Mateus Gustavo Chuqui: methodology, visualization, formal analysis. Frederico P. Brandini: project administration, writing - review & editing. Luigi Jovani: funding acquisition, writing - review & editing. Vivian H. Pellizari: supervision, writing - original draft, writing - review & editing.

## SUPPLEMENTARY MATERIAL

**Supplementary Table 1.** Environmental data, with the following information: ship station, sample number, latitude (LAT, °N), longitude (LONG, °E), bottom depth (BD, m), sampling depth (SD, m), temperature (T, °C), salinity (SAL, PSU), oxygen (O, mg/L), fluorescence (FLUOR, RFU), nitrite (NO_2-_, μM), nitrate (NO_3-_, μM), phosphate (PO_4_^3-^, μM), silicate (SiO^4-^, μM), and colored dissolved organic matter (CDOM, mg.L^−1^).

| Ship<br>Station | Sample | LAT<br>(°N) | LONG<br>(°E) | BD<br>(m) | SD<br>(m) | T<br>(°C) | SAL<br>(PSU) | O<br>(mg/L) | FLUOR<br>(RFU) | NO<sub>2</sub><sup>-</sup><br>(μM) | NO<sub>3</sub><sup>-</sup><br>(μM) | PO<sub>4</sub><sup>3-</sup><br>(μM) | SiO<sub>4</sub><sup>4-</sup><br>(μM) | CDOM<br>(mg.L<sup>-1</sup>) |
| --- | --- | --- | --- | --- | --- | --- | --- | --- | --- | --- | --- | --- | --- | --- |
| S0449 | 1 | -31.156 | -34.829 | 859 | 5 | 24.15 | 35.77 | 6.72 | 0.11 | 0.01 | 0.11 | 0.01 | 0.58 | 0.115 |
| S0449 | 2 | -31.156 | -34.829 | 859 | 51 | 22.42 | 35.85 | 6.96 | 0.13 | 0.01 | 0.06 | 0.01 | 0.5 | 0.222 |
| S0449 | 3 | -31.156 | -34.829 | 859 | 120 | 16.55 | 35.74 | 6.98 | 0.19 | 0.01 | NaN | NaN | NaN | 0.745 |
| S0449 | 4 | -31.156 | -34.829 | 859 | 400 | 11.85 | 35.07 | 6.52 | 0.17 | NaN | NaN | 0.32 | NaN | 1.047 |
| S0449 | 5 | -31.156 | -34.829 | 859 | 630 | 6.87 | 34.44 | 6.91 | 0.20 | NA | NA | NA | NA | 0.096 |
| S0449 | 6 | -31.156 | -34.829 | 859 | 800 | 5.58 | 34.32 | 7.18 | 0.22 | 0.01 | 29.5 | NaN | 13.91 | 1.009 |
| S0450 | 7 | -31.368 | -34.844 | 963 | 5 | 23.71 | 35.75 | 6.73 | 0.12 | 0.01 | 0.38 | NaN | 0.53 | 0.133 |
| S0450 | 8 | -31.368 | -34.844 | 963 | 10.8 | 23.66 | 35.75 | 6.76 | 0.12 | 0.01 | 0.01 | 0.01 | 0.53 | 0.132563 |
| S0450 | 9 | -31.368 | -34.844 | 963 | 32.5 | 22.30 | 35.80 | 7.05 | 0.13 | 0.01 | 0.06 | 0.01 | 0.49 | 0.1891 |
| S0450 | 10 | -31.368 | -34.844 | 963 | 43.5 | 20.75 | 35.91 | 7.39 | 0.14 | 0.01 | 0.13 | 0.01 | 0.48 | 0.224142 |
| S0450 | 11 | -31.368 | -34.844 | 963 | 53.1 | 20.48 | 35.93 | 7.42 | 0.14 | 0.01 | 0.15 | 0.01 | 0.72 | 0.256772 |
| S0450 | 12 | -31.368 | -34.844 | 963 | 62.2 | 19.98 | 35.92 | 7.41 | 0.15 | 0.01 | 0.05 | 0.01 | 0.73 | 0.295372 |
| S0450 | 13 | -31.368 | -34.844 | 963 | 76.0 | 18.75 | 35.90 | 7.41 | 0.16 | 0.01 | 0.06 | 0.01 | 0.55 | 0.377582 |
| S0450 | 14 | -31.368 | -34.844 | 963 | 94.0 | 18.39 | 35.90 | 7.33 | 0.18 | 0.01 | 0.14 | 0.02 | 0.72 | 0.525488 |
| S0450 | 15 | -31.368 | -34.844 | 963 | 105 | 17.51 | 35.83 | 7.06 | 0.19 | 0.01 | 0.11 | 0.03 | 1.04 | 0.615 |
| S0450 | 16 | -31.368 | -34.844 | 963 | 140 | 16.58 | 35.71 | 6.85 | 0.18 | 0.07 | 2.01 | 0.31 | 1.19 | 0.768 |
| S0450 | 17 | -31.368 | -34.844 | 963 | 450 | 11.73 | 35.04 | 6.48 | 0.17 | 0.01 | 17.99 | 1.44 | 5.38 | 1.031 |
| S0450 | 18 | -31.368 | -34.844 | 963 | 900 | 4.54 | 34.28 | 7.12 | 0.23 | 0.01 | 40.11 | NaN | 29.3 | 1.071 |
| S0452 | 19 | -31,204 | -34,761 | 701 | 7 | 23.38 | 35.83 | 6.57 | 0.13 | 0.01 | 0.15 | 0.01 | 0.6 | 0.165128 |
| S0452 | 20 | -31,204 | -34,761 | 701 | 125 | 17.39 | 35.78 | 7.24 | 0.21 | 0.06 | 0.87 | 0.26 | 0.81 | 0.662957 |
| S0452 | 21 | -31,204 | -34,761 | 701 | 450 | 11.02 | 34.94 | 6.47 | 0.19 | 0.01 | 16.71 | 1.4 | 4.89 | 1.029448 |
| S0452 | 22 | -31,204 | -34,761 | 701 | 610 | 7.46 | 34.51 | 6.77 | 0.21 | 0.01 | 32.58 | 0.56 | 13.79 | 1.048834 |
| S0468 | 23 | -30,891 | -35,716 | 1565 | 10.8 | 23.71 | 35.79 | 6.75 | 0.12 | 0.01 | 0.28 | 0.01 | 0.44 | 0.099546 |
| S0468 | 24 | -30,891 | -35,716 | 1565 | 100 | 17.83 | 35.83 | 7.29 | 0.22 | 0.03 | 0.64 | 0.09 | 0.64 | 0.617857 |
| S0468 | 25 | -30,891 | -35,716 | 1565 | 450 | 11.82 | 35.07 | 6.51 | 0.18 | 0.01 | 13.83 | NaN | 3.87 | 1.024779 |
| S0468 | 26 | -30,891 | -35,716 | 1565 | 850 | 4.65 | 34.27 | 7.17 | 0.23 | 0.01 | 37.41 | 0.92 | 24.8 | 1.0291 |
| S0468 | 27 | -30,891 | -35,716 | 1565 | 1400 | 3.01 | 34.56 | 5.97 | 0.26 | NaN | NaN | NaN | NaN | 1.598562 |
| S0470 | 28 | -30.669 | -35.774 | 819 | 5 | 24.26 | 35.72 | 6.06 | 0.14 | 0.01 | 0.4 | 0.01 | 0.59 | 0.124 |
| S0470 | 29 | -30.669 | -35.774 | 819 | 120 | 17.70 | 35.87 | 7.21 | 0.20 | 0.03 | 0.27 | 0.08 | 0.74 | 0.654 |
| S0470 | 30 | -30.669 | -35.774 | 819 | 450 | 10.97 | 34.95 | 6.52 | 0.19 | 0.01 | 20.69 | 0.91 | 6.19 | 1.018 |
| S0470 | 31 | -30.669 | -35.774 | 819 | 680 | 6.97 | 34.46 | 6.94 | 0.22 | 0.01 | NaN | 0.92 | 16.06 | 1.005 |
| S0472 | 32 | -30,781 | -35,963 | 1010 | 5 | 24.19 | 35.71 | 6.24 | 0.13 | 0.01 | 0.12 | 0.01 | 0.56 | 0.138258 |
| S0472 | 33 | -30,781 | -35,963 | 1010 | 90 | 17.62 | 35.83 | 7.22 | 0.21 | 0.11 | 0.25 | 0.09 | 0.65 | 0.656786 |
| S0472 | 34 | -30,781 | -35,963 | 1010 | 450 | 11.72 | 35.05 | 6.51 | 0.18 | 0.01 | 18.66 | 0.68 | 5.39 | 1.035621 |
| S0472 | 35 | -30,781 | -35,963 | 1010 | 950 | 4.19 | 34.30 | 6.89 | 0.23 | 0.01 | 42.94 | 0.88 | 33.6 | 1.135916 |
| S0474 | 36 | -30.869 | -35.956 | 1203 | 5 | 23.97 | 35.28 | 6.02 | 0.11 | 0.01 | 0.2 | 0.01 | 0.56 | 0.131 |
| S0474 | 37 | -30.869 | -35.956 | 1203 | 90 | 17.58 | 35.82 | 7.18 | 0.21 | 0.12 | 1.25 | 0.1 | 1.13 | 0.634 |
| S0474 | 38 | -30.869 | -35.956 | 1203 | 450 | 11.52 | 35.01 | 6.50 | 0.17 | 0.01 | 18.22 | 0.85 | 5.37 | 1.031 |
| S0474 | 39 | -30.869 | -35.956 | 1203 | 1100 | 3.52 | 34.35 | 6.45 | 0.25 | 0.01 | 36.97 | 0.81 | 36.69 | 1.319 |
| S0475 | 40 | -30,899 | -35,949 | 1246 | 0 | 23.93 | 35.46 | 6.35 | 0.12 | 0.01 | NaN | 0.01 | 0.4 | 0.16839 |
| S0475 | 41 | -30,899 | -35,949 | 1246 | 11.0 | 23.54 | 35.83 | 6.84 | 0.12 | 0.01 | 0.83 | 0.01 | 0.47 | 0.177183 |
| S0475 | 42 | -30,899 | -35,949 | 1246 | 33 | 22.88 | 35.91 | 7.02 | 0.12 | 0.01 | 0.83 | 0.01 | 0.48 | 0.189527 |
| S0475 | 43 | -30,899 | -35,949 | 1246 | 44 | 22.64 | 35.92 | 7.09 | 0.13 | 0.01 | 0.47 | 0.01 | 0.49 | 0.224445 |
| S0475 | 44 | -30,899 | -35,949 | 1246 | 54 | 19.97 | 35.91 | 7.45 | 0.14 | 0.01 | 0.28 | 0.01 | 0.48 | 0.285281 |
| S0475 | 45 | -30,899 | -35,949 | 1246 | 63 | 19.81 | 35.91 | 7.46 | 0.15 | 0.01 | 0.14 | 0.03 | 0.53 | 0.354657 |
| S0475 | 46 | -30,899 | -35,949 | 1246 | 77 | 19.15 | 35.91 | 7.45 | 0.17 | 0.01 | 0.37 | 0.05 | 0.56 | 0.445493 |
| S0475 | 47 | -30,899 | -35,949 | 1246 | 89 | 18.58 | 35.87 | 7.35 | 0.19 | 0.01 | 0.52 | 0.07 | 0.6 | 0.542412 |
| S0475 | 48 | -30,899 | -35,949 | 1246 | 107 | 17.26 | 35.79 | 7.15 | 0.20 | 0.1 | 1.4 | 0.13 | 0.78 | 0.676514 |
| S0475 | 49 | -30,899 | -35,949 | 1246 | 142 | 16.26 | 35.70 | 6.96 | 0.18 | 0.03 | 4.3 | 0.28 | 1.18 | 0.739518 |
| S0480 | 50 | -30.863 | -35.999 | 704 | 3 | 24.74 | 36.12 | 5.90 | 0.12 | 0.01 | 0.05 | 0.01 | 0.42 | 0.110 |
| S0480 | 51 | -30.863 | -35.999 | 704 | 115 | 17.62 | 35.83 | 7.23 | 0.23 | 0.08 | 0.59 | 0.07 | 0.5 | 0.655 |
| S0480 | 52 | -30.863 | -35.999 | 704 | 450 | 11.44 | 35.00 | 6.49 | 0.18 | 0.01 | 17.33 | 0.95 | 5.73 | 1.027 |
| S0480 | 53 | -30.863 | -35.999 | 704 | 650 | 6.61 | 34.41 | 7.08 | 0.21 | 0.01 | 27.97 | 1.31 | 13.74 | 0.982 |
| S0481 | 54 | -30,083 | -36,083 | 1807 | 8 | 24.47 | 35.83 | 6.66 | 0.12 | NaN | 1.78 | NaN | 0.75 | 0.128045 |
| S0481 | 55 | -30,083 | -36,083 | 1807 | 9.7 | 24.47 | 35.83 | 6.66 | 0.12 | 0.01 | 1.07 | 0.04 | NaN | 0.131626 |
| S0481 | 56 | -30,083 | -36,083 | 1807 | 29.3 | 23.12 | 35.90 | 7.00 | 0.12 | 0.01 | 0.12 | 0.01 | 0.48 | 0.177792 |
| S0481 | 57 | -30,083 | -36,083 | 1807 | 39.2 | 23.01 | 35.91 | 7.02 | 0.12 | 0.01 | 0.2 | 0.01 | 0.55 | 0.215827 |
| S0481 | 58 | -30,083 | -36,083 | 1807 | 47.9 | 21.93 | 35.89 | 7.11 | 0.13 | 0.01 | 0.13 | 0.02 | 0.46 | 0.275923 |
| S0481 | 59 | -30,083 | -36,083 | 1807 | 56.1 | 19.37 | 35.88 | 7.46 | 0.14 | 0.01 | 0.06 | 0.02 | 0.5 | 0.322131 |
| S0481 | 60 | -30,083 | -36,083 | 1807 | 68.6 | 19.21 | 35.89 | 7.49 | 0.15 | 0.01 | 0 | 0.01 | 0.45 | 0.403621 |
| S0481 | 61 | -30,083 | -36,083 | 1807 | 78.5 | 19.10 | 35.89 | 7.49 | 0.17 | 0.01 | 0.15 | 0.05 | 0.57 | 0.473846 |
| S0481 | 62 | -30,083 | -36,083 | 1807 | 95.4 | 18.14 | 35.84 | 7.40 | 0.20 | 0.04 | 0.35 | 0.1 | 0.52 | 0.587549 |
| S0481 | 63 | -30,083 | -36,083 | 1807 | 126.4 | 17.06 | 35.81 | 7.04 | 0.19 | 0.01 | NaN | NaN | NaN | 0.699559 |
| S0481 | 64 | -30,083 | -36,083 | 1807 | 450 | 11.20 | 34.98 | 6.53 | 0.18 | 0.01 | 20.49 | 1.26 | 6.22 | 1.055881 |
| S0481 | 65 | -30,083 | -36,083 | 1807 | 1100 | 3.42 | 34.37 | 6.35 | 0.25 | 0.01 | NaN | NaN | NaN | 1.345166 |
| S0481 | 66 | -30,083 | -36,083 | 1807 | 1400 | 2.98 | 34.56 | 5.96 | 0.26 | 0.01 | 42.59 | 1.07 | 53.83 | 1.62109 |
| S0481 | 67 | -30,083 | -36,083 | 1807 | 1610 | 2.96 | 34.63 | 6.13 | 0.26 | 0.01 | 39.63 | 1.11 | 53.96 | 1.702128 |
| S0485 | 68 | -31.046 | -36.199 | 748 | 5 | 24.08 | 35.73 | 6.64 | 0.12 | 0.01 | 0.17 | 0.01 | 0.47 | 0.145 |
| S0485 | 69 | -31.046 | -36.199 | 748 | 85 | 17.69 | 35.83 | 7.18 | 0.22 | 0.01 | 0.24 | 0.01 | 0.47 | 0.695 |
| S0485 | 70 | -31.046 | -36.199 | 748 | 450 | 11.66 | 35.04 | 6.52 | 0.18 | 0.01 | 16.24 | 0.73 | 4.97 | 1.023 |
| S0485 | 71 | -31.046 | -36.199 | 748 | 680 | 6.76 | 34.44 | 7.00 | 0.22 | 0.01 | 28.84 | 0.59 | 14.63 | 1.002 |
| S0500 | 72 | -30,936 | -36,262 | 826 | 10 | 24.19 | 35.79 | 6.70 | 0.12 | 0.01 | 0.13 | 0.01 | 0.68 | 0.159 |
| S0500 | 73 | -30,936 | -36,262 | 826 | 95 | 17.54 | 35.82 | 7.12 | 0.19 | 0.01 | NaN | NaN | NaN | 0.649084 |
| S0500 | 74 | -30,936 | -36,262 | 826 | 450 | 11.81 | 35.06 | 6.52 | 0.18 | NaN | NaN | NaN | NaN | 1.03605 |
| S0500 | 75 | -30,936 | -36,262 | 826 | 740 | 6.08 | 34.39 | 7.06 | 0.22 | 0.01 | 29.89 | 0.67 | 17.2 | 0.991455 |
| S0507 | 76 | -30.742 | -35.899 | 1449 | 5 | 23.86 | 35.72 | 6.71 | 0.12 | 0.01 | 0.1 | 0.01 | NaN | 0.178 |
| S0507 | 77 | -30.742 | -35.899 | 1449 | 120 | 17.29 | 35.78 | 7.10 | 0.18 | 0.03 | 0.26 | 0.01 | NaN | 0.662 |
| S0507 | 78 | -30.742 | -35.899 | 1449 | 450 | 11.57 | 35.02 | 6.48 | 0.19 | 0.01 | 17.35 | 0.7 | NaN | 1.044 |
| S0507 | 79 | -30.742 | -35.899 | 1449 | 800 | 5.45 | 34.32 | 7.18 | 0.22 | 0.01 | 31.19 | 0.59 | NaN | 0.988 |
| S0507 | 80 | -30.742 | -35.899 | 1449 | 1388 | 3.06 | 34.52 | 5.92 | 0.25 | 0.01 | 38.35 | 0.8 | NaN | 1.572 |

**Supplementary Table 2.** Results of principal component analysis (PCA) using environmental variables (temperature, salinity, oxygen, fluorescence, CDOM, phosphate, silicate, nitrite and nitrate). Principal component loadings >0.40 are shown in bold.

| **Variables** | **PC1** | **PC2** | **PC3** |
| --- | --- | --- | --- |
| Temperature | -0.401 | 0.221 | -0.187 |
| Salinity | -0.401 | 0.020 | -0.030 |
| Oxygen | -0.238 | -0.389 | 0.600 |
| Chlorophyll | -0.189 | -0.606 | 0.220 |
| Nitrite | -0.025 | -0.581 | -0.728 |
| Nitrate | 0.419 | 0.021 | 0.118 |
| Phosphate | 0.349 | -0.107 | -0.048 |
| Silicate | 0.377 | 0.025 | 0.039 |
| CDOM | 0.378 | -0.281 | 0.082 |
| **% Explanation** | **57.07** | **17.03** | **8.93** |

**Supplementary Table 3.** Cell abundance (cell.mL^−1^) of Synechococcus spp. (SYN), Prochlorococcus spp. (PRO), picoeukaryotes (PEUK), nanoeukaryotes (NANOEUK) and heterotrophic bacteria (HBAC) for each sample, considering the ship station, latitude (LAT, °N), longitude (LONG, °E), sampling depth (SD, m), and the corresponding water mass and pelagic zone.

| Ship station | Sample | LAT (°N) | LONG (°E) | SD (m) | SYN (cell.mL-1) | PRO (cell.mL-1) | PEUK (cell.mL-1) | NANOEUK (cell.mL-1) | HBAC (cell.mL-1) | Water mass | Pelagic zone |
| --- | --- | --- | --- | --- | --- | --- | --- | --- | --- | --- | --- |
| S0449 | 1 | -31.156 | -34.829 | 5 | 8.02E+02 | 2.76E+02 | 5.76E+02 | 8,43E+02 | 4,87E+08 | TW | EPI |
| S0449 | 2 | -31.156 | -34.829 | 51 | 2.92E+03 | 1.38E+02 | 6.49E+02 | 6,11E+01 | 8,52E+08 | TW | EPI |
| S0449 | 3 | -31.156 | -34.829 | 120 | 1.98E+02 | 3.55E+04 | 1.79E+03 | 1,80E+02 | 9,48E+08 | TW | EPI |
| S0449 | 4 | -31.156 | -34.829 | 400 | 2.52E+02 | 5.93E+01 | 1.85E+00 | 1,85E+01 | 2,12E+08 | SACW | MESO |
| S0449 | 5 | -31.156 | -34.829 | 800 | 1.96E+02 | 2.09E+02 | 2.22E+01 | 3,70E+01 | 1,78E+08 | AIW | MESO |
| S0450 | 6 | -31.368 | -34.844 | 5 | 4.88E+03 | 1.61E+02 | 6.78E+02 | 2,74E+02 | 5,39E+08 | TW | EPI |
| S0450 | 7 | -31.368 | -34.844 | 10.8 | 4.73E+03 | 1.44E+02 | 5.81E+02 | 5,00E+01 | 6,68E+08 | TW | EPI |
| S0450 | 8 | -31.368 | -34.844 | 32.5 | 3.65E+03 | 1.54E+02 | 9.57E+02 | 5,00E+01 | 7,86E+08 | TW | EPI |
| S0450 | 9 | -31.368 | -34.844 | 43.5 | 3.59E+03 | 1.91E+02 | 1.01E+03 | 7,04E+01 | 8,42E+08 | TW | EPI |
| S0450 | 10 | -31.368 | -34.844 | 53.1 | 3.37E+03 | 1.24E+02 | 8.48E+02 | 1,13E+02 | 6,52E+08 | TW | EPI |
| S0450 | 11 | -31.368 | -34.844 | 62.2 | 2.83E+03 | 1.50E+02 | 9.26E+02 | 7,22E+01 | 6,16E+08 | TW | EPI |
| S0450 | 12 | -31.368 | -34.844 | 76.0 | 2.02E+03 | 3.26E+02 | 5.80E+02 | 4,26E+01 | 4,56E+08 | TW | EPI |
| S0450 | 13 | -31.368 | -34.844 | 94.0 | 1.95E+03 | 3.12E+03 | 1.11E+03 | 9,07E+01 | 3,33E+08 | TW | EPI |
| S0450 | 14 | -31.368 | -34.844 | 105 | 4.81E+02 | 2.72E+04 | 2.03E+03 | 9,30E+02 | 4,62E+08 | TW | EPI |
| S0450 | 15 | -31.368 | -34.844 | 140 | 3.35E+02 | 2.59E+04 | 1.12E+03 | 7,22E+01 | 2,06E+08 | TW | EPI |
| S0450 | 16 | -31.368 | -34.844 | 450 | 1.69E+02 | 1.96E+02 | 1.67E+01 | 3,70E+01 | 9,58E+07 | SACW | MESO |
| S0450 | 17 | -31.368 | -34.844 | 900 | 5.46E+02 | 1.30E+02 | 2.41E+01 | 6,85E+01 | 4,96E+08 | AIW | MESO |
| S0452 | 18 | -31,204 | -34,761 | 7 | 2.93E+03 | 3.67E+02 | 4.57E+02 | 4,41E+02 | 2,24E+05 | TW | EPI |
| S0452 | 19 | -31,204 | -34,761 | 125 | 8.89E+02 | 3.99E+04 | 3.06E+03 | 4,24E+02 | 3,95E+05 | TW | EPI |
| S0452 | 20 | -31,204 | -34,761 | 450 | 4.74E+02 | 3.44E+02 | 2.59E+01 | 9,81E+01 | 1,19E+05 | SACW | MESO |
| S0452 | 21 | -31,204 | -34,761 | 610 | 4.98E+02 | 7.41E+01 | 1.11E+01 | 7,22E+01 | 9,46E+04 | AIW | MESO |
| S0468 | 22 | -30,891 | -35,716 | 10.8 | 3.10E+03 | 2.44E+02 | 6.11E+02 | 8,61E+02 | 1,57E+05 | TW | EPI |
| S0468 | 23 | -30,891 | -35,716 | 100 | 4.28E+02 | 4.07E+04 | 4.06E+03 | 6,04E+02 | 4,24E+05 | TW | EPI |
| S0468 | 24 | -30,891 | -35,716 | 450 | 1.65E+02 | 3.80E+02 | 5.00E+01 | 3,15E+01 | 5,33E+04 | SACW | MESO |
| S0468 | 25 | -30,891 | -35,716 | 850 | 1.63E+02 | 5.74E+01 | 0.00E+00 | 2,04E+01 | 5,61E+04 | AIW | MESO |
| S0468 | 26 | -30,891 | -35,716 | 1400 | 2.33E+02 | 2.59E+01 | 3.70E+00 | 1,48E+01 | 3,15E+04 | AIW | BATHY |
| S0470 | 27 | -30.669 | -35.774 | 5 | 3.81E+03 | 1.26E+02 | 6.09E+02 | 2,78E+01 | 2,01E+05 | TW | EPI |
| S0470 | 28 | -30.669 | -35.774 | 120 | 2.15E+02 | 2.98E+04 | 1.47E+03 | 2,96E+02 | 2,14E+05 | TW | EPI |
| S0470 | 29 | -30.669 | -35.774 | 450 | 1.63E+02 | 2.15E+02 | 1.67E+01 | 1,48E+01 | 4,97E+04 | SACW | MESO |
| S0470 | 30 | -30.669 | -35.774 | 680 | 9.44E+01 | 3.33E+01 | 1.85E+00 | 1,30E+01 | 7,71E+04 | AIW | MESO |
| S0472 | 31 | -30,781 | -35,963 | 5 | 2.50E+03 | 2.22E+02 | 7.26E+02 | 1,67E+01 | 7,22E+04 | TW | EPI |
| S0472 | 32 | -30,781 | -35,963 | 90 | 2.13E+02 | 3.81E+04 | 3.73E+03 | 3,22E+02 | 3,67E+05 | TW | EPI |
| S0472 | 33 | -30,781 | -35,963 | 450 | 9.63E+01 | 2.67E+02 | 3.89E+01 | 2,41E+01 | 5,24E+04 | SACW | MESO |
| S0472 | 34 | -30,781 | -35,963 | 950 | 1.41E+02 | 5.56E+01 | 2.04E+01 | 2,04E+01 | 4,77E+04 | AIW | MESO |
| S0474 | 35 | -30.869 | -35.956 | 5 | 1.12E+03 | 1.83E+02 | 6.26E+02 | 4,81E+01 | 1,31E+05 | TW | EPI |
| S0474 | 36 | -30.869 | -35.956 | 90 | 2.00E+02 | 3.33E+04 | 3.01E+03 | 2,96E+02 | 2,34E+05 | TW | EPI |
| S0474 | 37 | -30.869 | -35.956 | 450 | 1.30E+03 | 2.48E+02 | 1.85E+01 | 1,74E+02 | 6,87E+04 | SACW | MESO |
| S0474 | 38 | -30.869 | -35.956 | 1100 | 9.54E+02 | 5.00E+01 | 5.56E+00 | 8,15E+01 | 5,24E+04 | AIW | BATHY |
| S0475 | 39 | -30,899 | -35,949 | 5 | 4.40E+03 | 3.26E+02 | 6.87E+02 | 4,09E+02 | 4,05E+05 | TW | EPI |
| S0475 | 40 | -30,899 | -35,949 | 11.0 | 7.22E+03 | 1.89E+02 | 4.80E+02 | 5,41E+02 | 5,03E+05 | TW | EPI |
| S0475 | 41 | -30,899 | -35,949 | 33 | 3.63E+03 | 3.00E+02 | 7.22E+02 | 1,19E+02 | 4,54E+05 | TW | EPI |
| S0475 | 42 | -30,899 | -35,949 | 44 | 4.65E+03 | 4.78E+02 | 8.54E+02 | 2,15E+02 | 4,80E+05 | TW | EPI |
| S0475 | 43 | -30,899 | -35,949 | 54 | 3.34E+03 | 7.33E+02 | 1.13E+03 | 1,24E+02 | 4,56E+05 | TW | EPI |
| S0475 | 44 | -30,899 | -35,949 | 63 | 3.63E+03 | 1.33E+03 | 1.35E+03 | 2,50E+02 | 4,67E+05 | TW | EPI |
| S0475 | 45 | -30,899 | -35,949 | 77 | 2.40E+03 | 5.11E+03 | 2.63E+03 | 2,22E+02 | 5,65E+05 | TW | EPI |
| S0475 | 46 | -30,899 | -35,949 | 89 | 1.48E+03 | 2.26E+04 | 3.94E+03 | 8,52E+01 | 4,65E+05 | TW | EPI |
| S0475 | 47 | -30,899 | -35,949 | 107 | 1.47E+03 | 3.68E+04 | 2.89E+03 | 4,48E+02 | 4,03E+05 | TW | EPI |
| S0475 | 48 | -30,899 | -35,949 | 142 | 1.63E+03 | 1.02E+04 | 6.94E+02 | 2,44E+02 | 2,23E+05 | TW | EPI |
| S0480 | 49 | -30.863 | -35.999 | 3 | 1.53E+03 | 1.72E+02 | 7.06E+02 | 1,59E+02 | 2,96E+04 | TW | EPI |
| S0480 | 50 | -30.863 | -35.999 | 115 | 7.57E+02 | 4.30E+04 | 3.45E+03 | 2,13E+02 | 2,85E+05 | TW | EPI |
| S0480 | 51 | -30.863 | -35.999 | 450 | 2.81E+02 | 2.46E+02 | 2.59E+01 | 6,30E+01 | 7,13E+04 | SACW | MESO |
| S0480 | 52 | -30.863 | -35.999 | 650 | 3.67E+02 | 5.00E+01 | 1.85E+00 | 7,96E+01 | 5,93E+04 | AIW | MESO |
| S0481 | 53 | -30,083 | -36,083 | 8 | 3.69E+03 | 1.46E+02 | 7.98E+02 | 5,70E+02 | 1,29E+05 | TW | EPI |
| S0481 | 54 | -30,083 | -36,083 | 9.7 | 3.25E+03 | 1.17E+02 | 7.93E+02 | 7,59E+01 | 2,63E+05 | TW | EPI |
| S0481 | 55 | -30,083 | -36,083 | 29.3 | 2.24E+03 | 1.28E+02 | 9.61E+02 | 6,85E+01 | 4,06E+05 | TW | EPI |
| S0481 | 56 | -30,083 | -36,083 | 39.2 | 3.05E+03 | 1.67E+02 | 1.05E+03 | 1,19E+02 | 2,66E+05 | TW | EPI |
| S0481 | 57 | -30,083 | -36,083 | 47.9 | 2.97E+03 | 7.00E+02 | 1.68E+03 | 6,67E+01 | 2,88E+05 | TW | EPI |
| S0481 | 58 | -30,083 | -36,083 | 56.1 | 2.70E+03 | 5.57E+02 | 1.56E+03 | 1,76E+02 | 1,54E+05 | TW | EPI |
| S0481 | 59 | -30,083 | -36,083 | 68.6 | 2.32E+03 | 9.33E+02 | 2.26E+03 | 7,41E+01 | 2,41E+05 | TW | EPI |
| S0481 | 60 | -30,083 | -36,083 | 78.5 | 1.86E+03 | 1.22E+04 | 3.96E+03 | 3,70E+01 | 1,59E+05 | TW | EPI |
| S0481 | 61 | -30,083 | -36,083 | 95.4 | 1.66E+03 | 4.82E+04 | 4.68E+03 | 1,41E+02 | 1,58E+05 | TW | EPI |
| S0481 | 62 | -30,083 | -36,083 | 126.4 | 3.37E+02 | 3.66E+04 | 1.85E+03 | 1,30E+02 | 1,41E+05 | TW | EPI |
| S0481 | 63 | -30,083 | -36,083 | 450 | 3.22E+02 | 4.22E+02 | 4.63E+01 | 4,81E+01 | 4,92E+04 | SACW | MESO |
| S0481 | 64 | -30,083 | -36,083 | 1100 | 2.28E+02 | 8.33E+01 | 1.11E+01 | 2,41E+01 | 4,68E+04 | AIW | BATHY |
| S0481 | 65 | -30,083 | -36,083 | 1400 | 2.72E+02 | 3.33E+01 | 7.41E+00 | 3,89E+01 | 2,19E+04 | CSW | BATHY |
| S0481 | 66 | -30,083 | -36,083 | 1610 | 4.46E+02 | 3.52E+01 | 1.85E+00 | 2,59E+01 | 9,46E+04 | CSW | BATHY |
| S0485 | 67 | -31.046 | -36.199 | 5 | 4.47E+03 | 8.15E+01 | 7.80E+02 | 7,59E+01 | 1,25E+04 | TW | EPI |
| S0485 | 68 | -31.046 | -36.199 | 85 | 6.33E+02 | 5.12E+04 | 4.24E+03 | 6,67E+01 | 1,20E+05 | TW | EPI |
| S0485 | 69 | -31.046 | -36.199 | 450 | 3.80E+02 | 3.30E+02 | 5.19E+01 | 5,00E+01 | 3,35E+04 | SACW | MESO |
| S0485 | 70 | -31.046 | -36.199 | 680 | 2.24E+02 | 4.63E+01 | 3.70E+00 | 2,22E+01 | 1,80E+04 | AIW | MESO |
| S0500 | 71 | -30,936 | -36,262 | 10 | 3.68E+03 | 1.56E+02 | 8.78E+02 | 2,43E+02 | 3,51E+05 | TW | EPI |
| S0500 | 72 | -30,936 | -36,262 | 95 | 6.98E+02 | 4.19E+04 | 2.47E+03 | 1,65E+02 | 9,18E+05 | TW | EPI |
| S0500 | 73 | -30,936 | -36,262 | 450 | 1.31E+04 | 2.63E+02 | 1.48E+01 | 3,70E+01 | 1,70E+05 | SACW | MESO |
| S0500 | 74 | -30,936 | -36,262 | 740 | 4.28E+02 | 1.30E+01 | 3.70E+00 | 2,78E+01 | 1,26E+05 | AIW | MESO |
| S0507 | 75 | -30.742 | -35.899 | 5 | 4.77E+03 | 1.22E+02 | 7.57E+02 | 7,96E+01 | 2,35E+05 | TW | EPI |
| S0507 | 76 | -30.742 | -35.899 | 120 | 6.50E+02 | 3.23E+04 | 1.53E+03 | 1,26E+02 | 4,77E+05 | TW | EPI |
| S0507 | 77 | -30.742 | -35.899 | 800 | 3.15E+02 | 2.41E+01 | 7.41E+00 | 1,30E+01 | 1,33E+05 | SACW | MESO |
| S0507 | 78 | -30.742 | -35.899 | 450 | 3.94E+02 | 1.94E+02 | 1.67E+01 | 2,59E+01 | 1,42E+05 | AIW | MESO |
| S0507 | 79 | -30.742 | -35.899 | 1388 | 4.67E+02 | 7.41E+00 | 1.85E+00 | 2,78E+01 | 8,24E+04 | CSW | BATHY |

**Supplementary Table 4.** Means, standard deviations (SD), and minimum-maximum (min-max) range of picoplankton abundances (cell.mL^−1^) and relative abundance (%) from all stations and depths. Autotrophs: Synechococcus spp.; Prochlorococcus spp.; Picoeukaryotes. Heterotrophs: Heterotrophic bacteria.

| Picoplankton group | Abundance (cell.mL<sup>-1</sup>) |  | Range, N = 79 |  | Relative abundance (%) |  |
| --- | --- | --- | --- | --- | --- | --- |
|  | Mean | SD | Min | Max |  |  |
| *Synechococcus* spp. | 1.84E+03 | 2.06E+03 | 9.44E+01 | 1.31E+04 | 0.16 | 16.77 |
| *Prochlorococcus* spp. | 7.94E+03 | 1.50E+04 | 7.41E+00 | 5.12E+04 | 0.71 | 72.38 |
| Picoeukaryotes | 1.03E+03 | 1.22E+03 | 0.00E+00 | 4.68E+03 | 0.09 | 9.39 |
| Nano-eukaryotes | 1.60E+02 | 1.99E+02 | 1.30E+01 | 9.30E+02 | 0.01 | 1.46 |
| Heterotrophic bacteria | 1.12E+08 | 2.44E+08 | 1.25E+04 | 9.48E+08 | 99.02 | - |

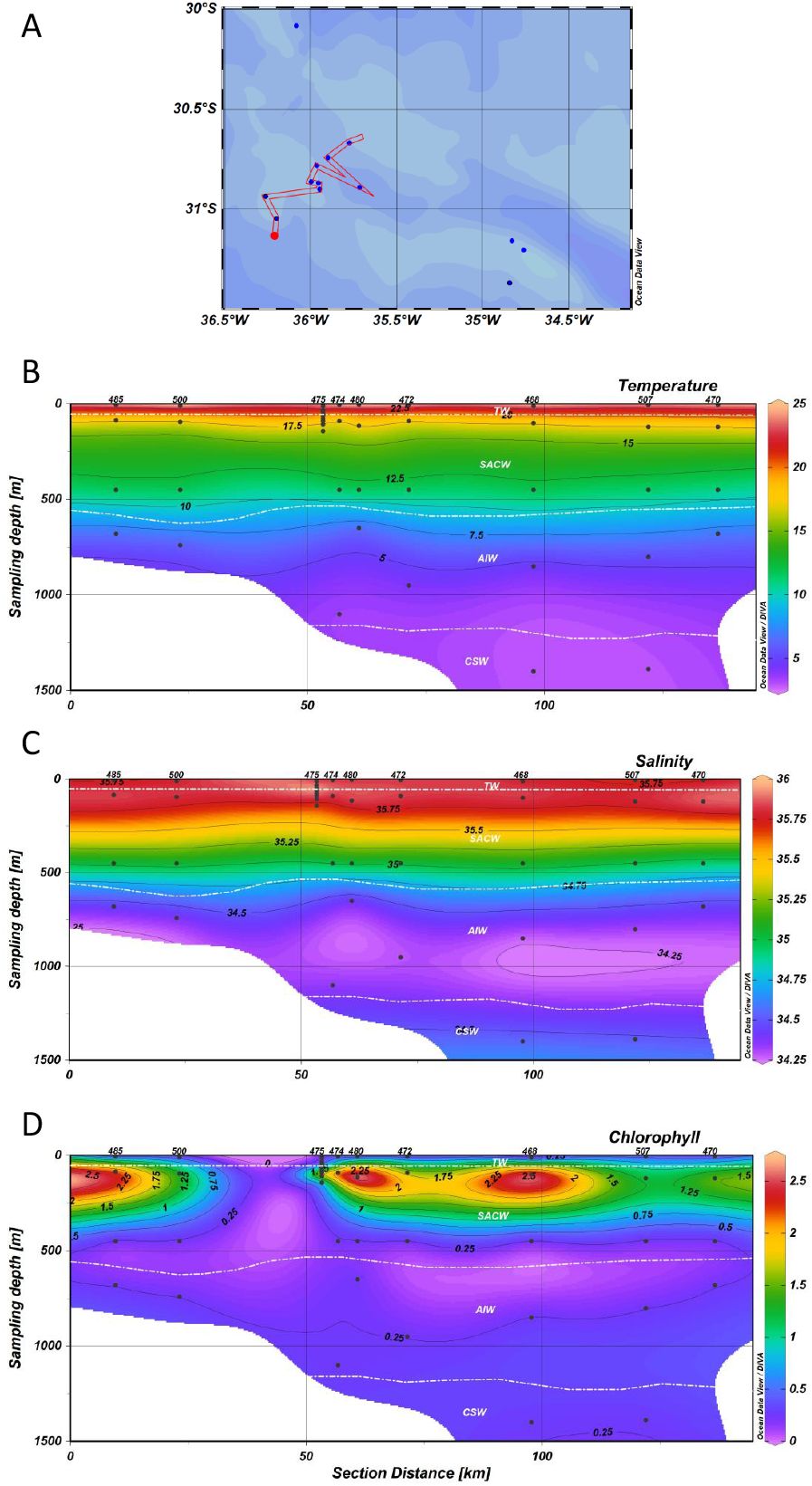

**Supplementary Table 5.** Results of PERMANOVA test with water masses (TW - Tropical Water, SACW - South Atlantic Central Water, AIW - Antarctic intermediate Water, and CSW - Circumpolar Superior Water) and pelagic zones (Epipelagic, Mesopelagic and Bathypelagic). Comparisons with p-value <0.05 are shown in bold.

| Group | p-value |
| --- | --- |
| Pelagic Zone | 0.001 |
| Epipelagic x Mesopelagic | 0.001 |
| Epipelagic x Bathypelagic | 0.008 |
| Mesopelagic x Bathypelagic | 0.589 |
| Water masses | > 0.5 |

**Supplementary Table 6.**
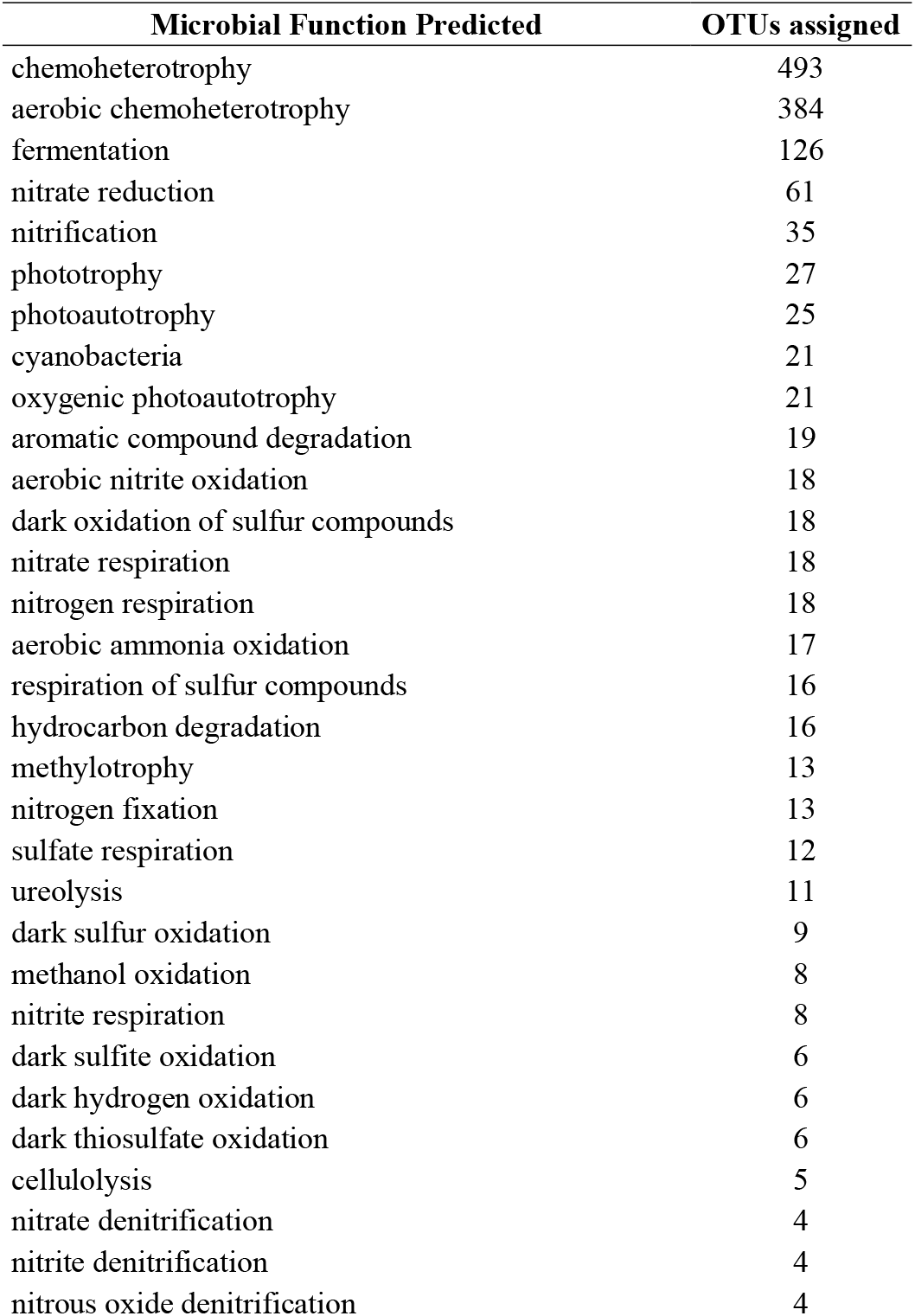
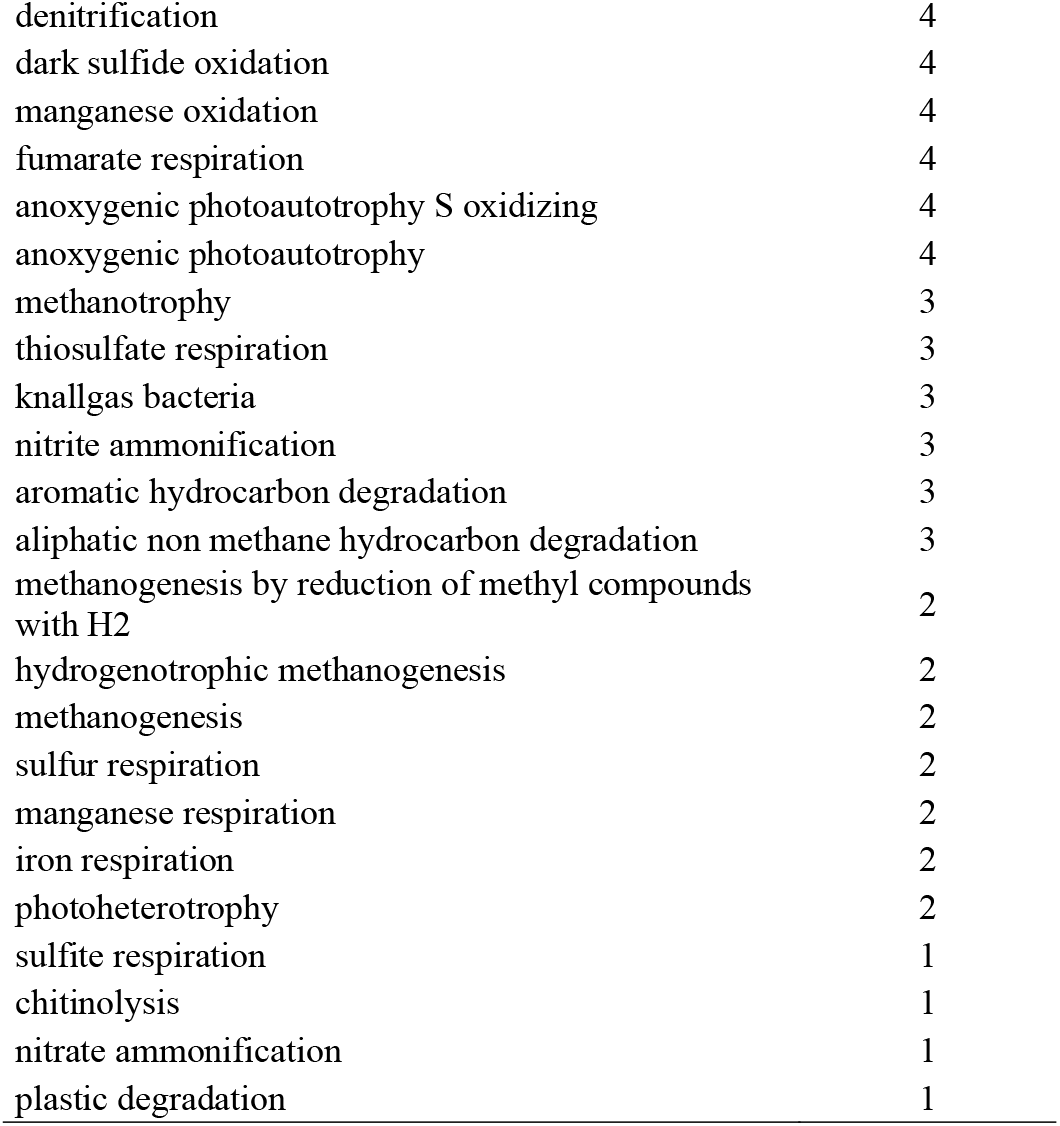
Microbial functional groups predicted in the pelagic zones from the RGR, and the number of bacterial and archaeal OTUs identified for each functional group.

| Microbial Function Predicted | OTUs assigned |
| --- | --- |
| chemoheterotrophy | 493 |
| aerobic chemoheterotrophy | 384 |
| fermentation | 126 |
| nitrate reduction | 61 |
| nitrification | 35 |
| phototrophy | 27 |
| photoautotrophy | 25 |
| cyanobacteria | 21 |
| oxygenic photoautotrophy | 21 |
| aromatic compound degradation | 19 |
| aerobic nitrite oxidation | 18 |
| dark oxidation of sulfur compounds | 18 |
| nitrate respiration | 18 |
| nitrogen respiration | 18 |
| aerobic ammonia oxidation | 17 |
| respiration of sulfur compounds | 16 |
| hydrocarbon degradation | 16 |
| methylotrophy | 13 |
| nitrogen fixation | 13 |
| sulfate respiration | 12 |
| ureolysis | 11 |
| dark sulfur oxidation | 9 |
| methanol oxidation | 8 |
| nitrite respiration | 8 |
| dark sulfite oxidation | 6 |
| dark hydrogen oxidation | 6 |
| dark thiosulfate oxidation | 6 |
| cellulolysis | 5 |
| nitrate denitrification | 4 |
| nitrite denitrification | 4 |
| nitrous oxide denitrification | 4 |
| denitrification | 4 |
| dark sulfide oxidation | 4 |
| manganese oxidation | 4 |
| fumarate respiration | 4 |
| anoxygenic photoautotrophy S oxidizing | 4 |
| anoxygenic photoautotrophy | 4 |
| methanotrophy | 3 |
| thiosulfate respiration | 3 |
| knallgas bacteria | 3 |
| nitrite ammonification | 3 |
| aromatic hydrocarbon degradation | 3 |
| aliphatic non methane hydrocarbon degradation | 3 |
| methanogenesis by reduction of methyl compounds<br>with H<sub>2</sub> | 2 |
| hydrogenotrophic methanogenesis | 2 |
| methanogenesis | 2 |
| sulfur respiration | 2 |
| manganese respiration | 2 |
| iron respiration | 2 |
| photoheterotrophy | 2 |
| sulfite respiration | 1 |
| chitinolysis | 1 |
| nitrate ammonification | 1 |
| plastic degradation | 1 |

**Supplementary Figure 1.**
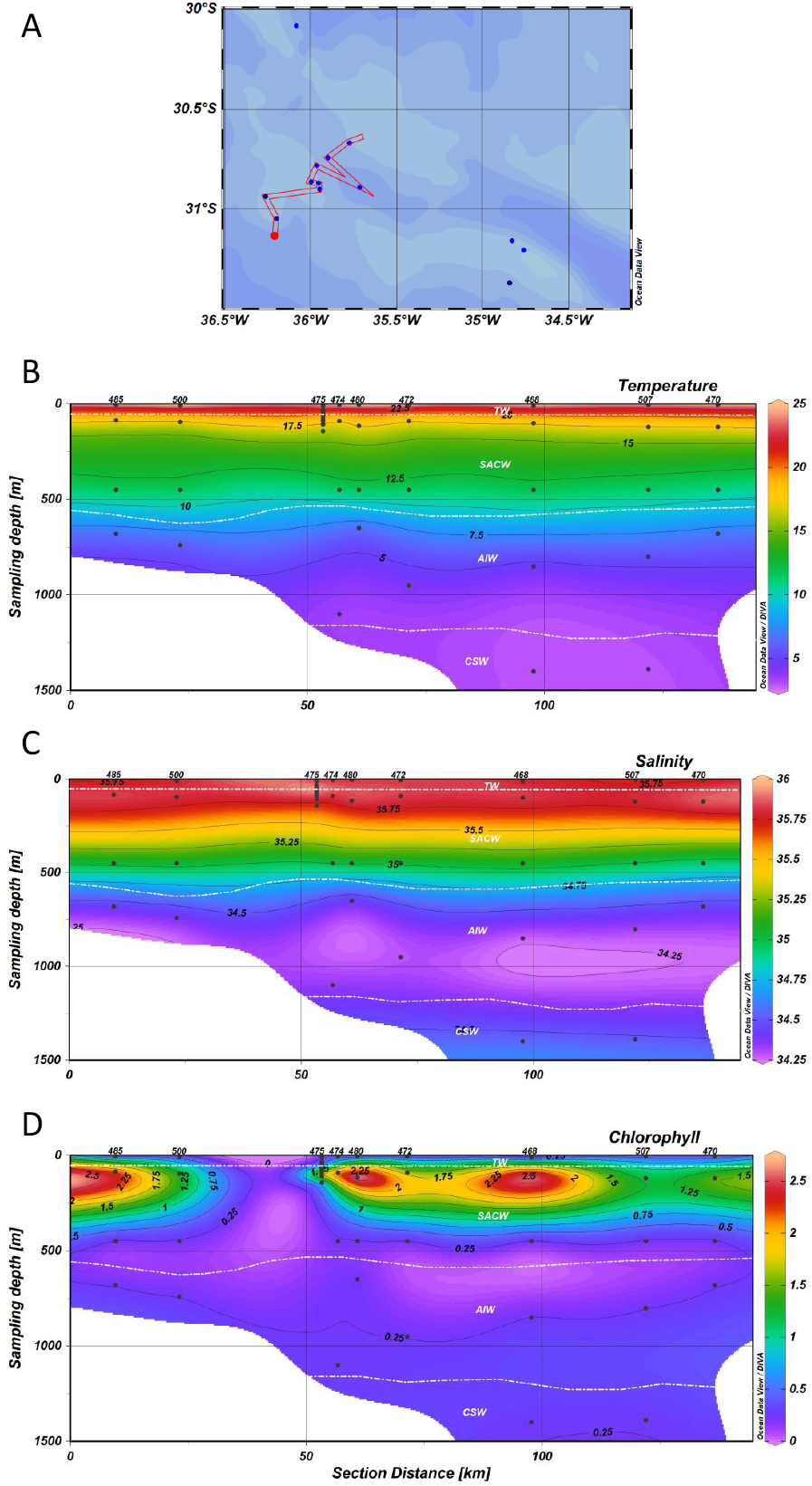
Depth profiles of hydrographic conditions to main sampling area in the Rio Grande Rise water column (0-1.500m): (A) West stations, (B) Temperature (°C), (C) Salinity (PSU) and (D) Chlorophyll (μg. L-^1^). Station labels are indicated at the top of each panel (west stations - 485, 500, 475, 474, 480, 472, 468, 507 and 470). Additional stations to the east were also used to statistics.

**Supplementary Figure 2.**
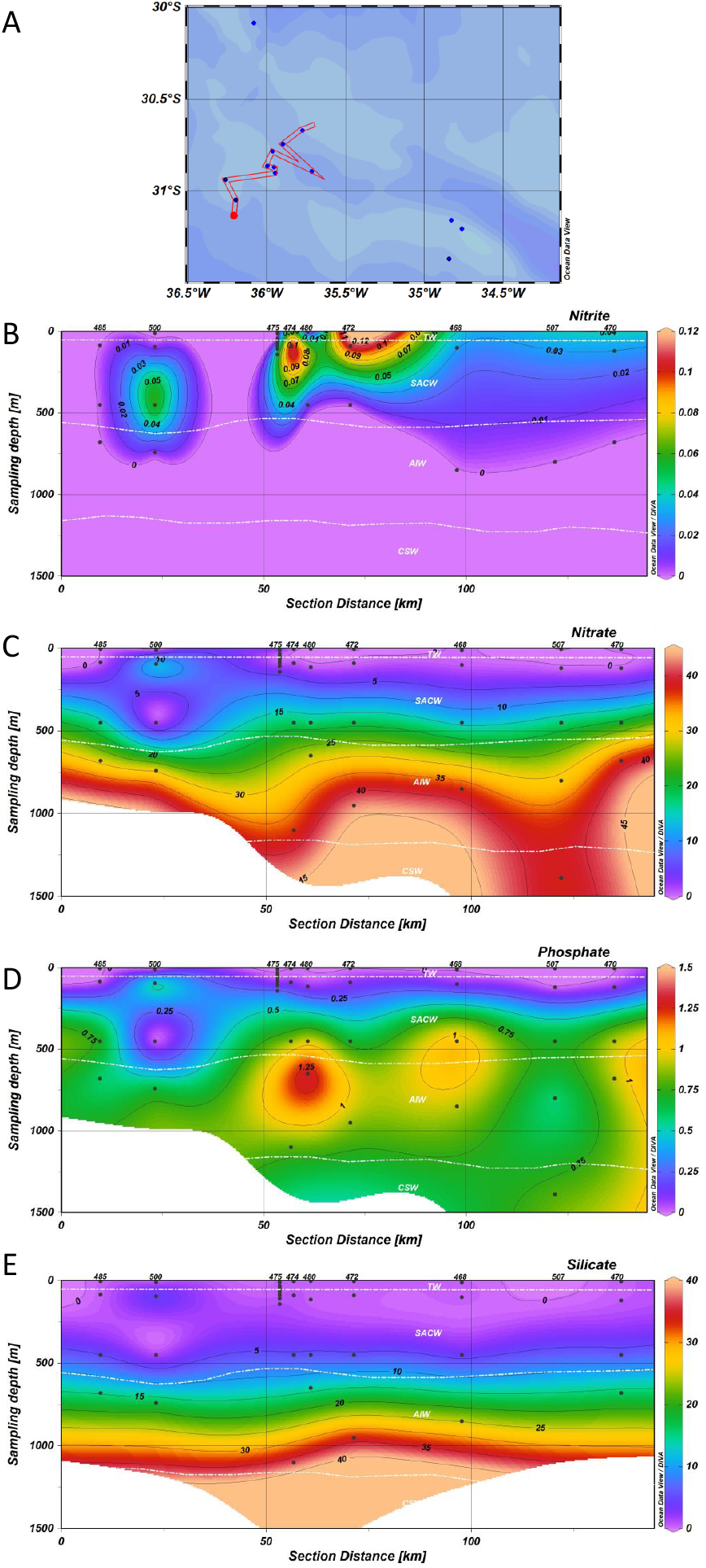
Concentrations of inorganic nutrients in the water column (0-1.500m) along the main sampling area in the Rio Grande: (A) West stations, (B) Nitrate (μM), (C) Nitrite (μM), (D) Phosphate (μM) and (E) Silicate (μM). Station labels are indicated at the top of each panel (west stations - 485, 500, 475, 474, 480, 472, 468, 507 and 470). Additional stations to the east were also used to statistics.

**Supplementary Figure 3.**
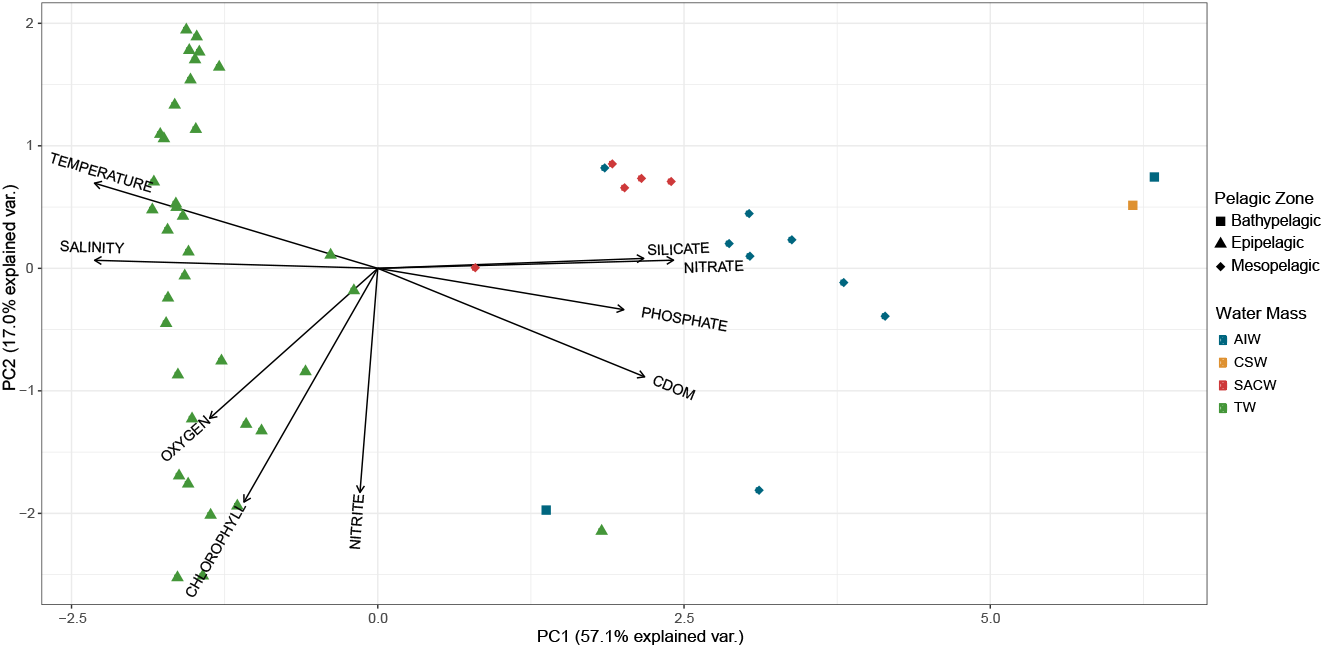
Oceanographic features of the study area. Outcome of the principal component analysis (PCA): Plot of principal components PC1 and PC2 showing the distribution of environmental variables (temperature, salinity, dissolved oxygen, colored dissolved organic matter (CDOM), fluorescence, phosphate, silicate, nitrite and nitrate. Circles represent samples collected on the epipelagic zone, squares represent samples collected in the mesopelagic zone, and triangles represent samples collected in the bathypelagic zone. Different colors represent the water masses: Red is Tropical Water (TW), blue is South Atlantic Central Water (SACW), green is Antarctic Intermediate Water (AIW) and yellow is Circumpolar Superior Water (CSW).

**Supplementary Figure 4.**
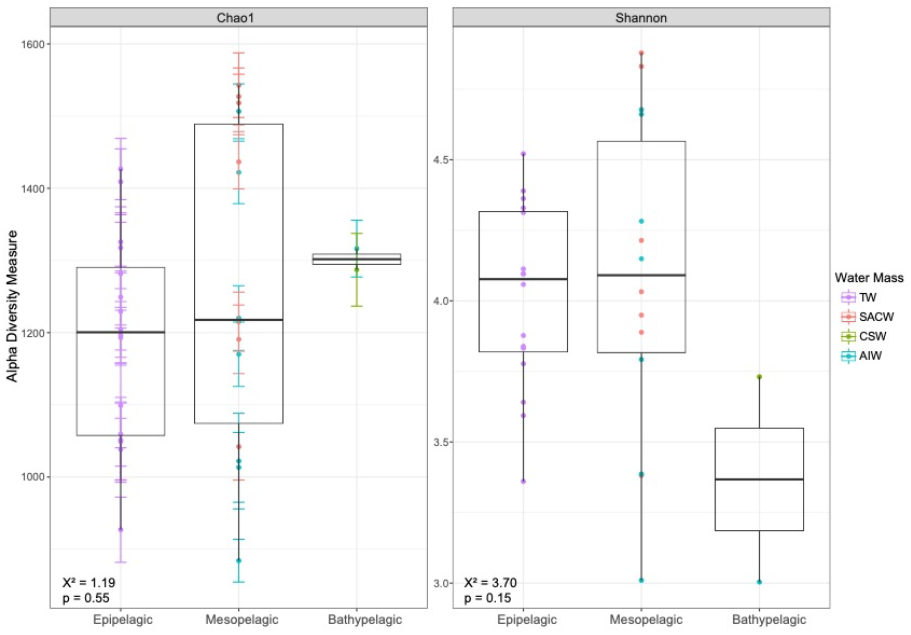
Alpha diversity (Shannon and Chao1 indexes) medians of microbial communities in pelagic zones in the RGR. Different colors represent the water masses: Purple is Tropical Water (TW), red is South Atlantic Central Water (SACW), blue is Antarctic Intermediate Water (AIW) and green is Circumpolar Superior Water (CSW).

**Supplementary Figure 5.**
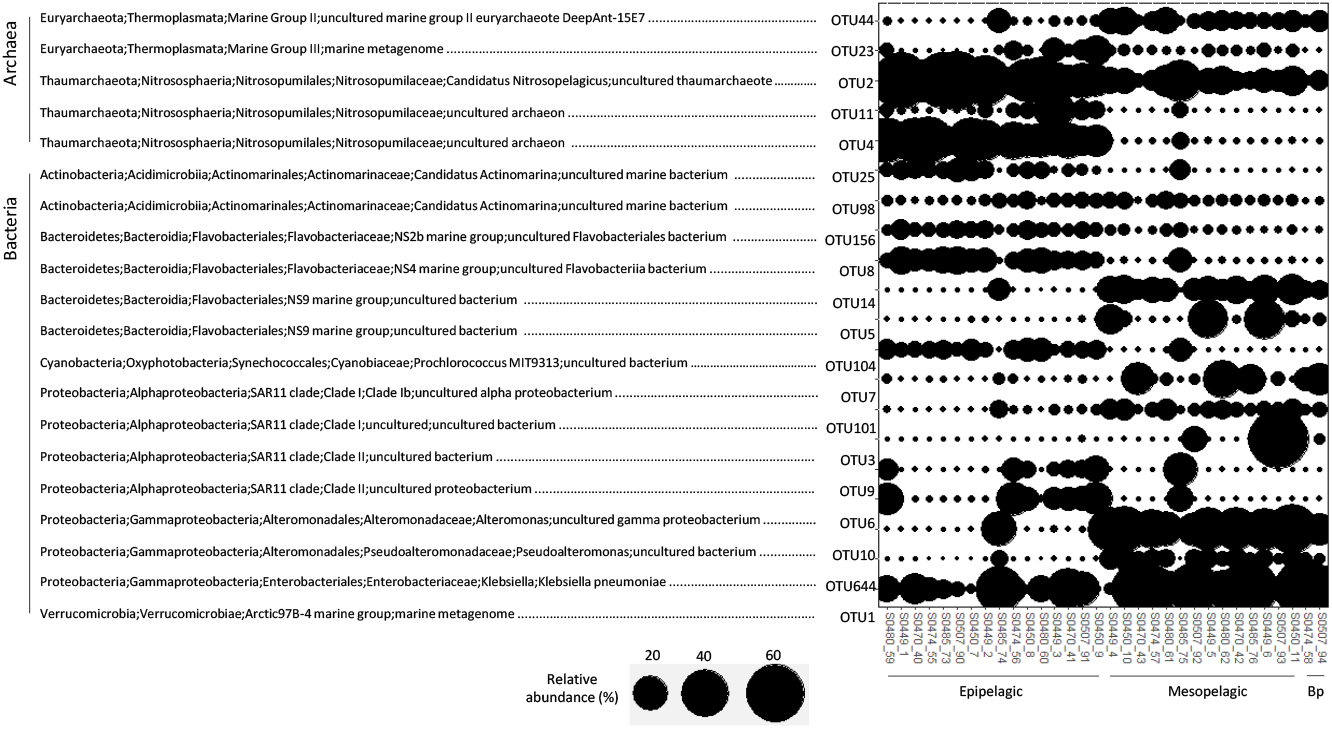
Classification of the most abundant OTUs for Archaea and Bacteria (> 0.1% relative abundance) in the pelagic zones. The size of circles is related to the relative abundance of each OTU. OTUs are organized by phylum. Black lines at the bottom indicate sample pelagic zone, i.e. epipelagic, mesopelagic and bathypelagic.

**Supplementary Figure 6.**
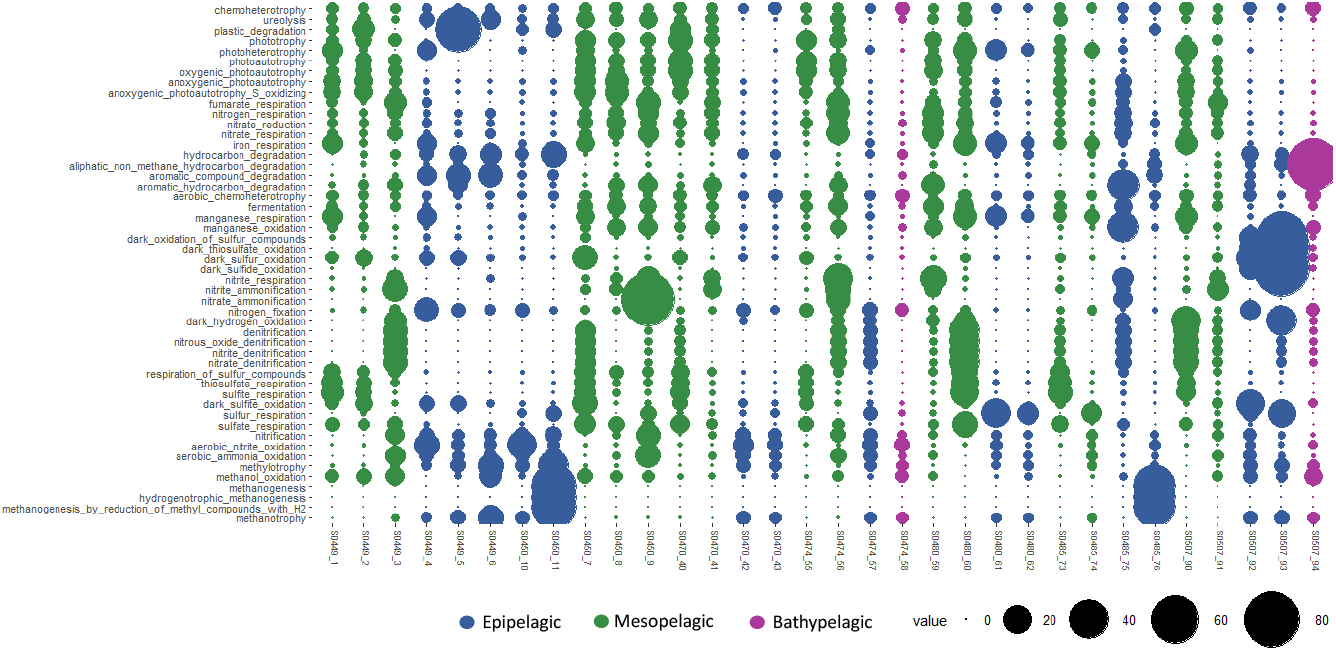
Mean of relative abundances of microbial functional groups for each pelagic zone of all sampling stations. Relative abundances are depicted in terms of circle size. Different colors represent the pelagic zones: blue is epipelagic, yellow is mesopelagic and green is bathypelagic.

## Notes

### Competing Interest Statement

The authors have declared no competing interest.

